# The Molecular Mechanism of Chloramphenicol and Thiamphenicol Resistance Mediated by a Novel Oxidase CmO in Sphingomonadaceae

**DOI:** 10.1101/2022.09.18.508450

**Authors:** Xiaodan Ma, Liying Zhang, Yijun Ren, Hui Yun, Hanlin Cui, Qian Li, Yuanqiang Guo, Shuhong Gao, Fengliang Zhang, Aijie Wang, Bin Liang

## Abstract

Antibiotic resistance mediated by bacterial enzyme inactivation plays a mysterious and crucial role for antibiotic degradation and selection pressure reduction in the environment. The enzymatic inactivation of the antibiotic chloramphenicol (CAP) involves nitro reduction, amide bond hydrolysis and acetylation modification. However, the molecular mechanism of enzymatic oxidation of CAP remains unknown. Here, a novel oxidase gene *cmO* was identified and confirmed biochemically to catalyze the resistance process through the oxidative inactivation at the side chain C-3’ position of CAP and thiamphenicol (TAP) in Sphingomonadaceae. The oxidase CmO is highly conservative in Sphingomonadaceae and shares the highest amino acid homology of 41.05% with the biochemically identified glucose methanol choline (GMC) oxidoreductases. Molecular docking and site-directed mutagenesis analyses demonstrated that CAP was anchored inside the protein pocket of CmO with the hydrogen bonding of key residues glycine (G)99, asparagine (N)518, methionine (M)474 and tyrosine (Y)380. CAP sensitivity test demonstrated that the acetyltransferase and CmO showed higher resistance to CAP as compared with the amide bond-hydrolyzing esterase and nitroreductase. This study provides a better theoretical basis and a novel diagnostic gene for understanding and assessing the fate and resistance risk of CAP and TAP in the environment.

**Importance:** Rising levels of antibiotic resistance undermines ecological and human health as a result of indiscriminate usage of antibiotics. Various resistance mechanisms have been revealed, for instance genes encoding proteins that degrade antibiotics, yet requiring further exploration. In this study, we reported a novel gene encoding an oxidase involved in the inactivation of typical amphenicol antibiotics (chloramphenicol and thiamphenicol), and the molecular mechanism was elucidated. The observation provides novel data to understand capabilities of bacteria to tackle antibiotic stress and suggests complex function of enzymes in the context of antibiotic resistance development and antibiotics removal. The reported gene can be further employed as an indicator to monitor amphenicols fate in the environment, benefiting the risk assessment in this era of antibiotic resistance.

## INTRODUCTION

The misuse and overuse of antibiotics, as the key driving force in the evolution and propagation of antibiotic resistance genes (ARGs), has caused ecological and human health risks, attracting the global attention (1-3). As the selection pressure posed by antibiotics was not strictly specific, long-term exposure to single or several antibiotics, especially those used widely and extensively, could accelerate the co-evolution of multiple ARGs (4-6). Amphenicol antibiotics, include chloramphenicol (CAP), its congener thiamphenicol (TAP) and florfenicol. CAP as the first potent antibiotic being commercialized globally to treat both human and livestock bacterial infections in quantities in the last 70 years (7). CAP is capable of preferentially interacting with the peptidyl transferase center (PTC) of 50S ribosomal subunit to restrain bacterial protein synthesis in Gram-positive/negative bacteria (8). Due to the hematological toxicity, CAP has been ceased in clinical use and live-stock farming in many economically developed countries (9-12). Despite all this, it remains active as the first-line prescription drug for the treatment of ophthalmic infections or other special multi-drug-resistant infections once again (13, 14). Importantly, CAP and TAP residues were still widely detected in various wastewater, soil, sediment, surface water and even some imported food at concentrations of nanograms to even milligrams per liter/kilogram, which especially applies to China (15-17). In consequence, the persistent selection pressure and (cross)induction from the ubiquitous CAP and TAP residues may lead to the evolution and transmission of ARGs in the environment, which brings a concealed hazard and risk to the ecosystems and human health. Currently, the biological indexes to indicate or predict CAP and TAP destruction process still lack and the relevant mechanisms are urgent to be uncovered.

A number of bacterial resistance mechanisms against CAP including efflux pump removal, ribosome protection and structural modification, and enzymatic inactivation, have been identified biochemically (13). Concretely, efflux pump membrane fusion protein encoding genes (e.g., *cmlA, cmlB, flo, floR, fexA, fexB*, and *pexA*) as a member of the main promoter superfamily (MFS), was not only the common regulatory gene of CAP efflux, but also associated with the other antibiotic resistance process. Gene *cfr* could encode RNA methyltransferase, and participate in the resistance response of various antibiotics, such as CAP (18-20). Bacterial enzyme inactivation plays a mysterious but as yet crucial role in the antibiotic degradation and selection pressure reduction in the environment (21-24). Although the enzymatic inactivation of CAP is one way of the bacterial antibiotic resistance, diverse metabolic processes via well-annotated functions (CAP acetyltransferase, nitroreductase, and amido bond hydrolase) could weaken the co/cross-selection pressure of CAP for environmental microbial communities. Bacterial CAP acetyltransferase is responsible for reversibly acetylating the hydroxyl group at the side chain C-1’ or C-3’ position of CAP (25-27). An esterase family hydrolase encoded by gene *estDL136* was biochemically confirmed to hydrolyze the amide bond of CAP and florfenicol side chain simultaneously (28). Moreover, a nitroreductase encoded by the gene *nfsB* can achieve rapid nitro reduction of CAP to amino-CAP (13, 29).

Very recently, *Sphingobium* sp. CAP-1 and *Sphingomonas* sp. CL5.1 isolated from activated sludge were reported to oxidize the C-3’ position linking with a hydroxyl group in CAP side chain to produce O-CAP. O-CAP can be further transformed into *p*-nitrobenzoic acid (PNBA) and mineralized through the protocatechuic acid (PCA) pathway(16, 30). The transformation of amphenicol antibiotics (CAP or TAP) to O-CAP or O-TAP is a novel oxidation-mediated detoxication process (16, 31). However, the key oxidase genes involved in this oxidative pathway need to be biochemically verified, and the corresponding molecular mechanism remains unknown.

Here, a novel CAP oxidase gene *cmO* from the glucose-methanol-choline (GMC) oxidoreductase family that catalyzes the oxidation of CAP into O-CAP in Sphingomonadaceae (*Sphingobium* sp. CAP-1 and *Sphingopyxis* sp. GC21), was successfully cloned and expressed in *Escherichia coli*, and biochemically characterized. The resistance characteristics of CmO were further compared with those of the reported CAP-degradative resistance genes encoding enzymatic nitro group reduction, amide bond hydrolysis, and acetylation. Finally, the key catalytic amino acid residues of CmO were identified, and the corresponding molecular mechanism was revealed. As the first report on CAP and TAP resistance mediated by an oxidase, the present study provides a better theoretical basis and a novel diagnostic gene for understanding and assessing the fate and resistance risk of CAP and TAP in the environment.

## MATERIALS AND METHODS

### Chemicals

CAP and TAP (both purity >99%) were purchased from Macklin (Shanghai, China). Other antibiotics and chemical reagents used were of the highest analytical purity.

### Bacterial strains, plasmids, primers and culture conditions

The bacterial strains, plasmids and the primers used in this study are listed in Table S1 and S2. *Sphingobium* sp. CAP-1 was grown at 30°C in Luria–Bertani (LB) medium or mineral salt medium (MSM, pH 7.0), each supplemented with CAP or TAP(16). *Escherichia coli* strains were grown at 37°C in LB medium, in which, antibiotics ampicillin (Amp, 100 mg/L), gentamicin (Gm, 100 mg/L), spectinomycin (Sm, 100 mg/L), streptomycin (Str, 100 mg/L) and kanamycin (Km, 50 mg/L) were added as required.

### Screening out a novel CAP oxidase in strain CAP-1

The comparative proteomic analysis of strain CAP-1 was previously carried out through isobaric tags for relative and absolute quantitation (iTRAQ) technology (16). Based on the genomic annotation and the fold change (FC) of the upregulated-expression proteins, one candidate gene was screened out (FC > 1.5 based, *P* < 0.05 on Student’s t-test, and the associated false discovery rate within 0.05 based on the Benjamin algorithm). The gene was further amplified with FastPfu DNA polymerase from genomic DNA of strain CAP-1 with the primer pair of ChoF/R (Table S2). The PCR products were ligated into pMD18-T vector and transformed into *E. coli* DH5α(32). The positive transformants were applied for the biotransformation assay. The oxidative biotransformation of CAP or TAP to O-CAP or O-TAP was confirmed using HPLC and high-resolution mass spectrometry as described below. A multiple alignment of the amino acid sequence of CmO and other biochemically identified oxidases (dehydrogenases) was performed as previously described (33, 34). Phylogenetic analysis of CmO and homologous oxidases was performed by using MEGA (version 7.0).

### Overexpression and purification of CmO

The *cmO* gene from the genomic DNA of strain CAP-1 was amplified with primers *cmO*-F/*cmO*-R (Table S2). The *cmO* fragment was ligated into a NdeI/XhoI-digested pET29a(+) plasmid using the ClonExpress II one-step cloning kit (Vazyme Biotech, China) to produce pET29a-*cmO*. The recombinant plasmid was subsequently transformed into *E. coli* BL21(DE3) and the positive transformant was cultured in LB medium containing Km (50 mg/L) at 37°C until the OD_600_ reached 0.6. Protein expression was induced at 16°C with 0.8 mM IPTG for 12 h. The cells of *E. coli* BL21 harboring pET29a-*cmO* were collected after centrifuging at 15,000g and 4°C for 20 min and washing twice with phosphate buffered saline (PBS, pH 7.4). The suspended cells were sonicated (Ultrasonic Cell Crusher, Thermo Fisher Scientific) and centrifuged under the conditions above to remove the cell debris. CmO was purified by using a Ni-nitrilotriacetic acid (Ni-NTA)-Sefinose column (Sangon Biotech, Shanghai, China) and dialyzed in Tris-HCl buffer (pH 7.4) overnight to remove imidazole and Ni^2+^. The purity and molecular mass of CmO were shown by SDS-PAGE, and the protein concentration was determined with the UV absorption peak at 280 nm by Nanodrop One (Thermo Scientific, USA).

### Enzyme assay

The enzyme activity was measured in 1 mL PBS (10 mM, pH=7.4) supplemented with the proper amount of purified CmO at 30°C for 10-20 min in triplicate. CAP and TAP were used as the reaction substrates with various initial concentrations, separately. Each enzymatic reaction was terminated by boiling for 1 min, and diluted by the equal volume methanol. Controls with inactive enzyme were also carried out under the same conditions. The determination of kinetic parameters including *V*_max_ and *K*_m_ were described elsewhere (35).

To determine the optimum temperature of CmO, the relative activities were compared at 10-80°C. In different buffers (36), the optimum pH was determined by comparing the enzyme activities in the pH range from 3.8 to 10.6. To determine the stability, CmO was pretreated in different temperature and pH for 1 h to analyze the remaining activity, respectively. The effects of metal ions, inhibitors and detergents on the enzyme activity were analyzed as previously described (35). The concentration of all metal ions and inhibitors were 1 mM. The detergents that comprised Triton X-100, Tween-20, Tween-80, sodium dodecyl sulfate (SDS) and cetyltriethyl ammonium bromide (CTAB) were added at a final concentration of 0.1% (m/V).

### Homology modeling and molecular docking

The 3D structure of the target protein, CmO (UniProtKB AC: A0A6I5YGT3), was constructed using an online protein structure prediction server, SWISS-MODEL (http://swissmodel.expasy.org/). The template protein was identified through BLAST sequence search. The molecular dynamics (MD) simulations were performed in Yinfo Cloud Computing Platform (YCCP, https://cloud.yinfotek.com/) to optimize the protein structure. The quality of protein model was evaluated using the online website SAVES V6 (https://saves.mbi.ucla.edu/)(35).

Molecular docking was conducted in Yinfo Cloud Computing Platform. By the DOCK 6.9 program, CAP was docked into the enzyme CmO pocket. Based on the quality of the protein, the optimum binding mode was selected for analysis (37-39).

### Site-directed mutagenesis

The mutations were introduced into CmO (expressed on pET-29a(+)) using a Mut Express II Fast Mutagenesis Kit (Vazyme, Nanjing, China) according to the manufacturer’s protocols. The primers used for site-directed mutagenesis of CmO were shown in Table S2, and the mutant amino acid residues included glycine (G)99, tyrosine (Y)380, methionine (M)474 and asparagine (N)518.

### Expression of CmO in *Sphingomonas wittichii* RW1

The fragment of *cmO* was amplified from the genomic DNA of strain CAP-1 with the primers pBBcmoF/R (Table S2). The PCR product was then ligated into KpnI/EcoRI-digested broad-host-range plasmid pBBR1-MCS2 using the ClonExpress II one-step cloning kit (Vazyme Biotech, China) to generate pBBR-*cmO*, which was electroporated into *S. wittichii* RW1 (a non-CAP degrader), and selected on LB plates supplemented with Sm (100 mg/L) and Km (50 mg/L).

### Analytical methods

CAP, TAP, O-CAP and O-TAP were analyzed using a HPLC (model-1260, Agilent Technologies, USA) equipped with a C_18_ column (4.6×100 mm, 2.7 μm, Agilent Technologies, USA). The mobile phase was a mixture of acetonitrile and ultrapure water containing 0.1% formic acid (30:70; vol/vol) at a flow rate of 0.6 mL/min in isocratic mode. A UV detector (model-1260, Agilent Technologies, USA) was set at 245 nm and 275 nm. For the refined identification of the O-TAP and O-CAP chemical structure, UPLC-triple-quadrupole mass spectrometer (UPLC-TQMS) was operated with appropriate parameters mentioned elsewhere (40). The high-resolution mass spectrometry and nuclear magnetic resonance (NMR) analysis using a 700 MHz liquid NMR spectrometer (AVANCE III, Bruker Biospin, Switzerland) were employed as described previously (16). The sample for NMR was separated by a high performance preparative liquid chromatography, freeze-dried to a solid powder and locked with deuterated chloroform (CDCl_3_) (16, 41, 42).

### Calculation of IC_50_ toward to CAP with different CAP resistant strains

Three types of CAP resistance via enzymatic inactivation including an amide bond-hydrolyzing esterase gene *estDL136* (28), a nitroreductase gene *nfsB* (13), and a acetyltransferase gene *cat* (43), were commercially synthesized and cloned into the multiple clone sites of *Nde*I/*Xho*I in vector pET-29a(+) (GenScript, USA), and then were transformed into *E. coli* BL21(DE3) to generate BL29dl136, BL29nfsB, and BL29cat, respectively. Microbroth serial dilution assays were employed to determine IC_50_ (half maximal inhibitory concentration) values for CAP and in a modification of EUCAST standard protocol. 100 μL volumes of Mueller-Hinton broth (MHB) were added into 96-well plates with CAP concentrations ranging from 0.125 to 256 μg/mL. The above constructed strains and BL29cmO were cultured overnight in MHB with Km and inoculated into fresh MHB to the OD_600_ value of 0.1. Diluted cultures were added to the prepared CAP-containing 96-well plates in 100 μL aliquots for an initial OD_600_ and volume of 0.05 and 200 μL, respectively. Growth with 6 biological replicates in the microplate constant temperature shaker at 800 rpm for 36 h at 30°C was measured by a Spark 10M multimode microplate reader (Tecan, Switzerland). Dose-response curves and significant differences in IC_50_ were analyzed and calculated as described elsewhere (13).

## RESULTS AND DISCUSSION

### Identification and functional verification of a novel oxidase gene *cmO*

The CAP resistance process in strain CAP-1 mainly relied on the oxidation of C-3’ position linking with a hydroxymethyl group of CAP to form the corresponding carboxyl group. The genes involved in this process (i.e. the genes being significantly up-regulated) were screened out based on the proteomic data (16). Specifically, an oxidase gene (annotated GL174_01150, GenBank accession number OP019282) that up-regulated significantly (3.197-fold) was screened out to verify its capacity to oxidize CAP. Based on the BLASTP analysis, the open reading frame (ORF) containing 1617 bp shared homology with many biochemically characterized GMC oxidoreductases. This ORF along with 200 bp upstream sequence was subcloned into the vector pMD18-T and transformed into *E. coli* DH5α. A positive transformant named DH18cmo was further screened out to verify its oxidizing capacity, which could completely oxidize the added CAP/TAP (160 μM) to O-CAP/O-TAP within 30 h based on the UPLC-TQMS and NMR analysis (Figure 1A). Moreover, two different metabolites were detected in the CAP and TAP biotransformation sample based on the total ions chromatograph (TIC) under the full-scan MS2 mode, separately (Figure 1B). The peak of CAP end-product mainly contained ions with *m/z* values of 108.2, 147.9, 192.2 and 334.9 [M − H]^−^, which was consistent with the ion fragment fingerprint determined for the O-CAP identification in a previous study (Figure 1C)(16). The peak from TAP metabolism was presumed to be O-TAP with *m/z* values of 108.1, 148.1, 184.1, and 368.1 [M − H]^−^. The freeze-dried samples were analyzed simultaneously using the NMR to ensure accurate chemical structure of this product.

**Figure 1.**
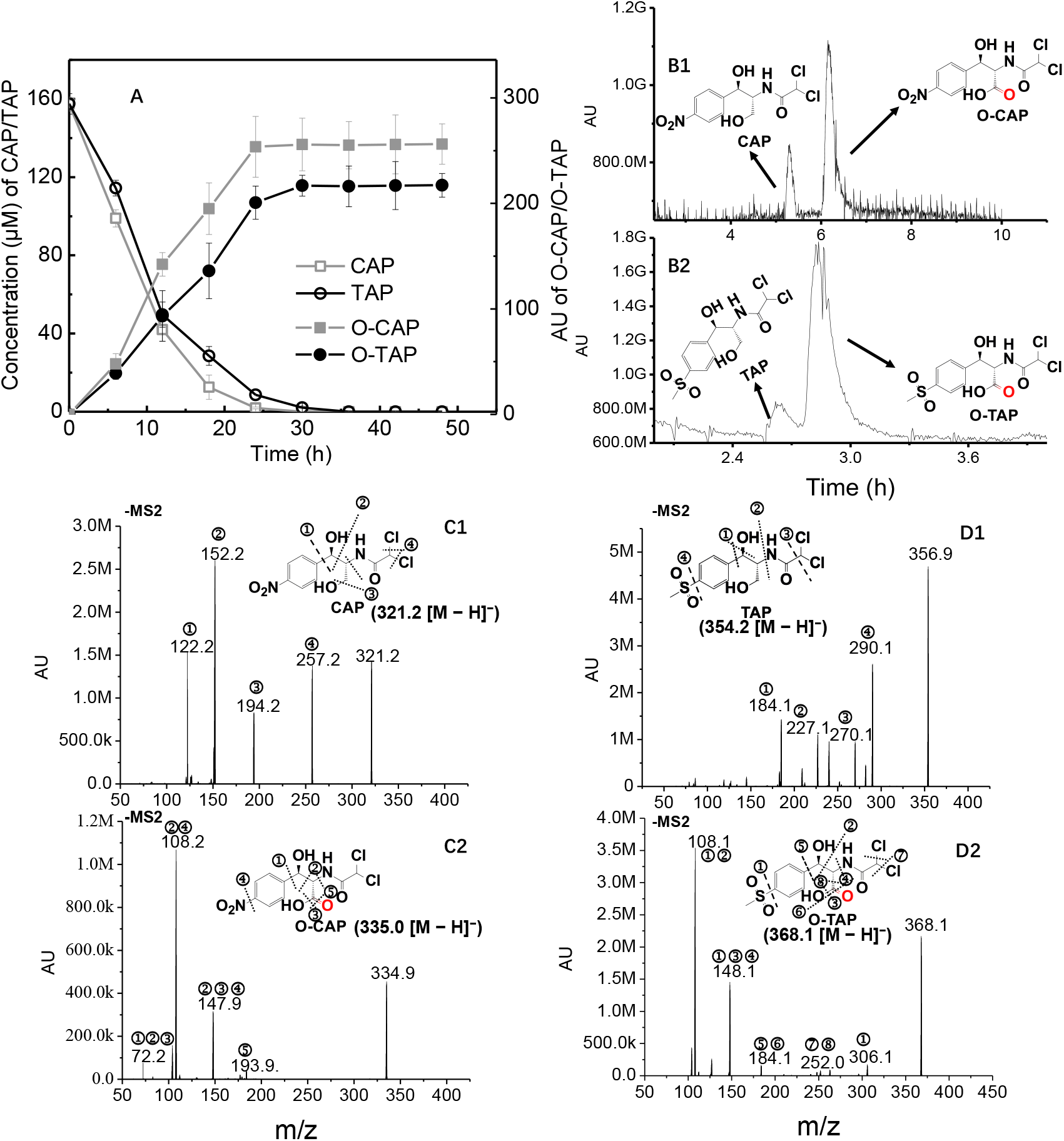
Transformation of CAP and TAP (160 μM) to O-CAP and O-TAP, respectively, by the expressed CmO in *E. coli* (DH18cmo) (A). Total ion chromatography (TIC) of antibiotic substrates (CAP and TAP) and oxidized products (O-CAP and O-TAP) in the full-scan MS2 mode (B1 and B2). HPLC-MS2 spectra of CAP (C1), O-CAP (C2), TAP (D1), and O-TAP (D2).

The product obtained was a white powder and the molecular formula was determined to be C_12_H_13_Cl_2_NO_6_S based on the high-resolution electrospray ionization mass spectrometry (HR-ESIMS) ion at *m/z* 367.9765 [M – H] ^−^ (calculated for C_12_H_12_Cl_2_NO_6_S, 367.9762) and the NMR data (Table 1). To elucidate its structure, 1D and 2D NMR experiments were performed. Three proton signals [*δ*_H_ 6.31 (1H, s), 5.43 (1H, d, *J* = 2.1 Hz), and 4.56 (1H, dd, *J* = 2.1 Hz)] were displayed from the ^1^H NMR spectrum (Figure S1A), besides a set of *p*-substituted benzene protons [*δ*_H_ 7.86 (2H, d, *J* = 8.2 Hz) and 7.67 (2H, d, *J* = 8.2 Hz)] in the low-field. Additionally, a methyl singlet [*δ*_H_ 3.07 (3H, s)] was also detected from the ^1^H NMR spectrum. Corresponding to the benzene ring displayed in the ^1^H NMR spectrum, 6 olefinic carbons [*δ*_C_ 150.0, 140.6, 128.5×2, and 128.0×2] assignable to a *p*-substituted benzene ring were observed from the ^13^C NMR spectrum (Figure S1B). Apart from these olefinic carbon signals, 2 carbonyls (*δ*_C_ 175.4 and 165.7), 3 methine carbons (*δ*_C_ 74.0, 67.7, and 61.4), and a methyl (*δ*_C_ 44.5) were also revealed according to the ^13^C, DEPT, and HMQC spectra (Figures S1CD). The ^1^H and ^13^C NMR spectra resembled those of the product O-CAP, for which the structure was elucidated and reported previously(16). After using the HMQC spectrum (Figure S1D) to assign the protons to the corresponding carbons (Table 1), the analysis of ^1^H–^1^H COSY and the HMBC spectra revealed the presence of 2-carboxy-2-(2,2-dichloroacetamido)-1-hydroxyethyl moiety, which was the same as that of O-CAP (Figure S1EF). The subsequent HMBC data analysis showed the correlations of H-1’ [*δ*_H_ 5.43 (1H, d, *J* = 2.1 Hz)] with C-1 (*δ*_C_ 150.0) and C-2 (*δ*_C_ 128.0), and H-2/6 [7.67 (2H, d, *J* = 8.2 Hz)] with C-1’ (*δ*_C_ 74.0), which suggested that the *p*-substituted benzene ring was attached at C-1’ of the 2-carboxy-2-(2,2-dichloroacetamido)-1-hydroxyethyl moiety. The methylsulfonyl group was deduced to be linked at C-4, which was supported by the HR-ESIMS (Figure S1G) and the chemical shift of C-4 (*δ*_C_ 140.6). There were no chemical reactions or changes in the chiral carbons during the transformation of TAP to O-TAP, and the configurations of C-1’ and C-2’ in O-TAP were the same as those of TAP. Thus, O-TAP was finally elucidated as (1’*S*,2’*S*)-2’-(2”,2”-dichloroacetamido)-3’-hydroxy-3’-(4(methylsulfonyl)phenyl)propanoic acid (Figure 2).

**Table 1.** NMR Data for Compound O-TAP. (*δ* in ppm, 100 MHz for ^13^C and 400 MHz for ^1^H, in CD_3_OD)^*a*^

| Position | $\delta_{\text{C}}$ | $\delta_{\text{H}}$ |
| --- | --- | --- |
| 1 | 150.0, C |  |
| 2 | 128.0, CH | 7.67 (2H, d, $J = 8.2$ Hz) |
| 3 | 128.5, CH | 7.86 (2H, d, $J = 8.2$ Hz) |
| 4 | 140.6, C |  |
| 5 | 128.5, CH | 7.86 (2H, d, $J = 8.2$ Hz) |
| 6 | 128.2, CH | 7.67 (1H, d, $J = 8.2$ Hz) |
| 1' | 74.0, CH | 5.43 (1H, d, $J = 2.1$ Hz) |
| 2' | 61.4, CH | 4.56 (1H, d, $J = 2.1$ Hz) |
| 3' | 175.4, C |  |
| 1'' | 165.7, C |  |
| 2'' | 67.7, CH | 6.31 (1H, s) |
| -SO<sub>2</sub>CH<sub>3</sub> | 44.5, CH<sub>3</sub> | 3.07 (3H, s) |

<sup>a</sup>Assignments of NMR data are based on $^1\text{H}$ , $^{13}\text{C}$ , DEPT, $^1\text{H}$ - $^1\text{H}$ COSY, HSQC, and HMBC experiments.

**Figure 2.**
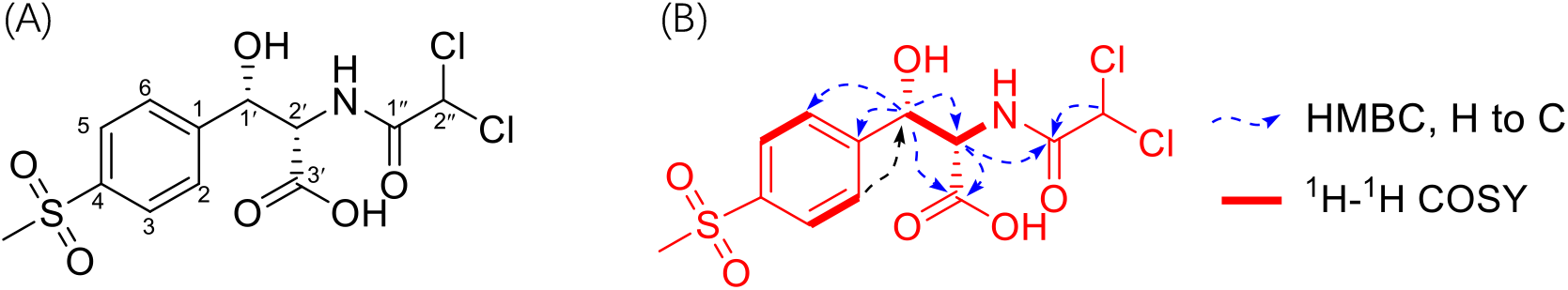
The chemical structure of O-TAP (A) and the ^1^H-^1^H COSY and key HMBC correlations in O-TAP (B).

A previous study reported that *Sphingomonas* sp. CL5.1 oxidized the C-3’ position with a hydroxymethyl group in TAP to O-TAP, yet only based on the high-resolution mass spectrometry analysis(31). This mode of oxidation is similar to our previously identified oxidation mode of CAP, of which O-CAP was the oxidative product based on the NMR analysis(16). Here, for the first time, we provided a NMR evidence to prove that the C-3’ position of TAP was also oxidized. Furthermore, the 1617 bp ORF was confirmed to be involved in the oxidation of the C-3’ position with a hydroxymethyl group in CAP/TAP to the corresponding carboxyl group and thus designated as *cmO* (chloramphenicol oxidase gene).

The recombinant strain RW1-*cmO* was capable of transforming 66.78±4.42 μM CAP to O-CAP by the expression of plasmid pBBR-*cmO*, and the *S. wittichii* RW1 control stain remained sensitive to CAP (Figure S2). Obviously, *cmO* can confer the CAP resistance to this type strain from Sphingomonadaceae. In addition, it was indirectly proved that *cmO* has the potential to rescue other sensitive bacteria at similar taxonomic status by mediating CAP resistance.

### CAP catabolic pathway initiated by CmO in Sphingobacteria

CmO is a novel oxidase that relieves the environmental stress caused by CAP and TAP, and can benefit the management of ecological risks of amphenicol antibiotics. A gene cluster comprising *cmO* and *pnbAB* (encoding transformation of PNBA to PCA) responsible for the upstream CAP-catabolic pathway was found highly conserved in *Sphingobium* sp. CAP-1, *Sphingopyxis* sp. GC21 and *Sphingomonas* sp. CL5.1 from Sphingomonadaceae (Figure 3AB). The gene *cmO* in strain CAP-1 was found to be similar to the oxidases from *Sphingopyxis* sp. GC21 and *Sphingomonas* sp. CL5.1 (98.83-98.88% identities). Phylogenetic analysis showed that CmO shared low homology (38.94 to 41.05%) with other biochemically characterized GMC family oxidoreductases (available in NCBI Swiss-Prot protein database), especially oxygen-dependent choline dehydrogenase BetA (Figure 3C). The oxidases from GMC family can oxidize a wide range of aliphatic and aromatic aldehydes or alcohols to the corresponding carboxylic acids, using NAD(P)^+^ or FAD as an electron acceptor (44, 45).

**Figure 3.**
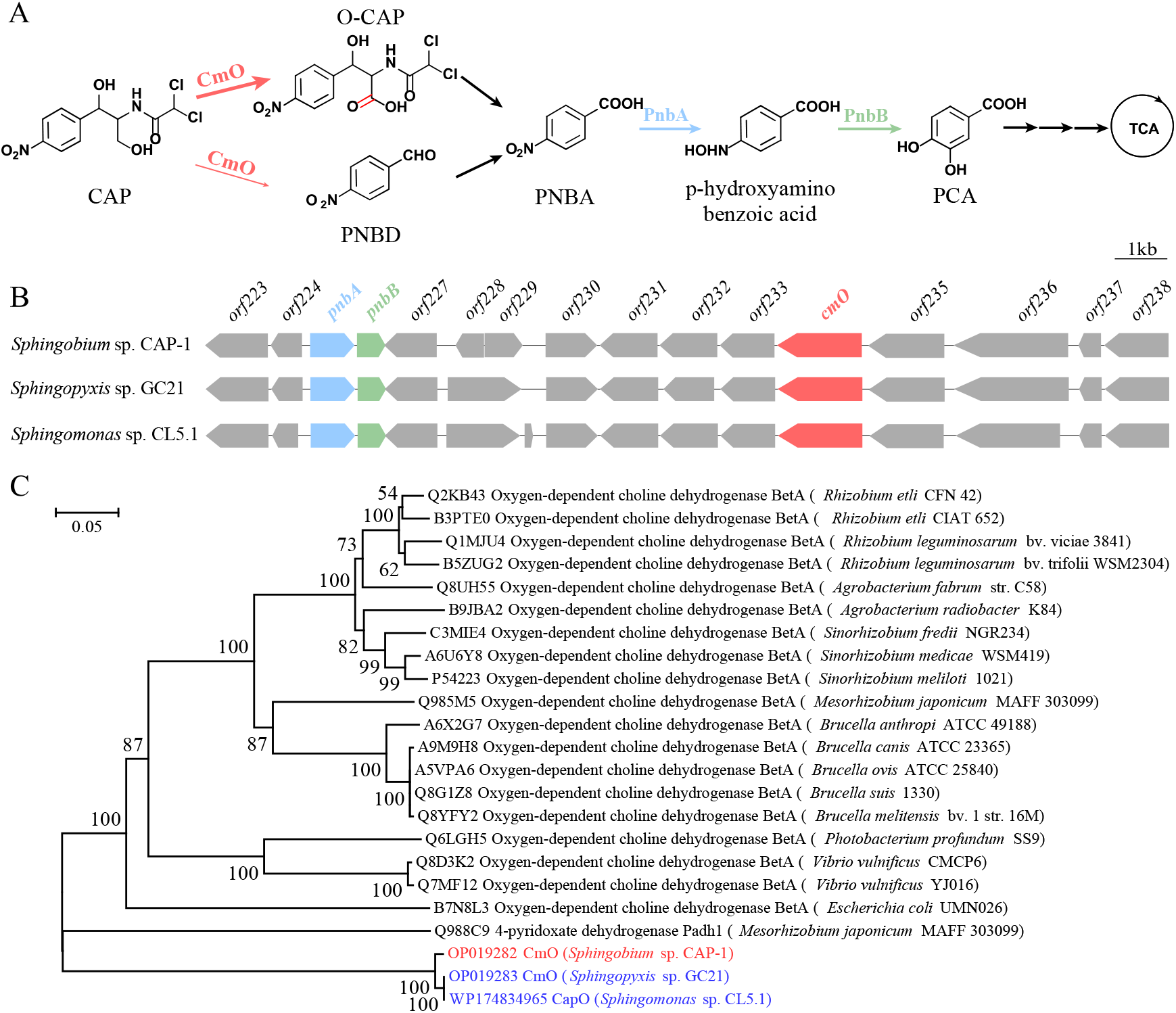
Candidate genes involved in CAP degradation in three strains of Sphingomonadaceae (*Sphingobium* sp. CAP-1, *Sphingopyxis* sp. GC21, and *Sphingomonas* sp. CL5.1). The proposed CAP catabolic pathway: the pink thick arrow indicates the major catalytic reaction mediated by CmO, and the pink thin arrows indicates the minor catalytic reaction; O-CAP, the oxidation product of the C-3’ position in CAP benzene side chain; PNBD, *p*-nitrobenzaldehyde; PNBA, *p*-nitrobenzoic acid, and PCA, protocatechuate (A). Genetic determinants of the upstream catabolism of CAP including *cmO* and *pnbAB* (B). Phylogenetic analysis of CmO revealing its relationship to other GMC oxidoreductase family members (C).

### Biochemical characterization of CmO

The *cmO* fragment was cloned into the vector pET-29a(+) and transformed into *E. coli* BL21(DE3). One positive transformant named BL29cmo was selected to verify its oxidative ability. O-CAP and O-TAP was continually accumulated during the oxidative degradation from CAP and TAP (35 μM) within 24-30 h, separately (Figure. S3). After the optimization experiment, the expressed protein was purified from the crude extract by using the Ni-nitrilotriacetic acid affinity chromatography with 0.8 mM IPTG induction. The purified, denatured protein showed a single band on SDS-PAGE, with a molecular mass slightly lower than 66.4 kDa, which was in agreement with its theoretical molecular mass of 59.26 kDa (Figure S4). CmO can catalyze CAP to produce O-CAP, *p*-nitrobenzaldehyde (PNBD) and PNBA (Figure S5A). Based on the UPLC-based analysis, approximate 100 μM CAP was completely transformed to 82.90%±3.09% O-CAP and 17.10 %±0.78% PNBD, and 44.64%±0.52% of PNBD was further oxidized to PNBA (Figure S5B). CmO could also oxidize PNBD to PNBA directly (Figure S5C), but could not further transform O-CAP (Figure S5D). Taken together, the resistance process mediated by CmO mainly comprises the oxidation of CAP to O-CAP (the major catalytic reaction) and PNBD (the minor catalytic reaction), and the further weak oxidation of PNBD to PNBA, as shown in Figure 3A. The effect of FAD, FMN, NAD^+^ and NADP^+^ on catalytic efficiency was compared with that of electron acceptor-free (CK, 100%), and FAD contributed to the maximum relative activity of CmO (144.99%±2.53%), as shown in Figure S6. The similar catalytic characteristics were observed when substituting TAP for CAP as the initial substrate.

The biochemical characterization of CmO was performed by investigating the effects of temperature, pH values, metal ions, and chemical inhibitors/detergents on the catalytic activity and stability. CmO was capable of keeping the stable catalytic activity in the range of 10-40°C, and gradually lost its stability once exceeding 50°C. The optimum temperature was 30°C for catalytic reaction process *in vitro* (Figure S7A). CmO exhibited the highest catalytic ability at pH 7.4. With the acidification and alkalization of the buffer systems, the relative activity of CmO finally decreased to 18.52±3.00% (pH=3.8) and 10.09%±0.55% (pH=10.6), respectively. CmO could maintain the catalytic stability between pH 7.0 and 7.4 for 1 h preincubation, and became unstable during the acidification or alkalization process (Figure S7B).

The effect of various metal ions on CmO catalytic activity was different under the optimum condition. 1 mM Co^2+^, Cr^3+^, Cd^2+^, Hg^2+^ and Ag^+^ strongly inhibited CmO catalytic activity (reduced 83.74-96.28% relative activity), whereas 1 mM Cu^2+^, Fe^2+^, Ni^2+^ and Mn^2+^ decreased 28.81-45.85% activity of CmO. Interestingly, 1 mM Zn^2+^ and Mg^2+^ had a promoting effect on the enzyme catalytic process, increasing the catalytic activity by of 14.63±7.35% and 22.11±4.10%, respectively. Ca^2+^, Na^+^, Fe^3+^, K^+^, Mo^2+^ and Al^3+^ had no significant influence on CmO activity (Figure S7C). All the tested chemical inhibitors and detergents showed the varying degrees of inhibition on CmO. Incubation of CmO with 1 mM iodide acetyl, nitrilotriacetic acid (NTA), phenyl-methane sulfonyl fluoride (PMSF), diethyl pyrocarbonate (DEPC) and 0.1% hexadecyl trimethyl ammonium bromide (CTAB, *m/v*) for 1 h obviously inhibited its catalytic activity, reducing 85.23-98.30% activity. CmO lost 15.96-33.27% catalytic activity with the addition of ethylene diamine tetraacetie acid (EDTA), 1,10-phenolene and 2,2’-bipyridine. The detergents, namely sodium dodecyl sulfate (SDS), TritonX-100, Tween-20 and Tween-80 at 0.1% (*m/v*) level, decreased the activity of CmO within 25.07-59.55% (Figure S7D). Importantly, CmO remained 66.73±3.14% activity after the inhibition by EDTA addition, which indicated that it was not a metal ion dependent oxidase.

In the 1 mL enzymatic reaction system, the initial catalytic rate *V*_0_ of CmO was detected and calculated at 50, 100, 200, 300, 400, 500 and 700 μM CAP and TAP, respectively. The kinetics of the CmO-mediated oxidation of CAP and TAP were apparent; specifically, 459 μg/L purified CmO with a 300 μM FAD cofactor for the oxidation of CAP and TAP possessed a *V*_max_ of 199.22±3.44 and 177.59±3.54 μM/s, *K*_m_ of 70.36±6.68 and 72.36± 4.58μM, *k*_cat_ of 140.28±2.42 and 137.83±2.75 s^-1^, and catalytic efficiency (*k*_cat_/*K*_m_) of 1.99 and 1.90 μM^-1^ s^-1^ separately, based on the Hill equation (n=1, the analogical Michaelis-Menten equation).

### The molecular mechanism of CmO mediated oxidative catalysis

Multiple alignment of the amino acid sequence of CmO with several biochemically characterized GMC oxidoreductases revealed the highly conserved amino acid residue sites (G99, Y380, M474, and N518) that were potentially related to the key catalytic activities of CmO homologs from the CAP-degrading bacteria in Sphingomonadaceae (Figure S8). However, these amino acid residues differed in other biochemically characterized GMC oxidases. In oxygen-dependent choline dehydrogenases, the Y380 and M474 residues were both replaced by the alanine (A) residue, G99 and N518 residues remained constant (46-50). The glutamate (E) 312 residue was the key binding site of choline (46), which was substituted by the aspartic acid (D) residue in CmO. Moreover, the valine (V) 464 and histidine (H) 466 residues were involved in the oxidation process of choline to betaine aldehyde(46). The G99 residue was highly conserved in most GMC family members and played a vital role in the anchoring of FAD (51). PhcC, PhcD, CHD, and HMFO belonging to this family are able to oxidize the hydroxyl group connected with benzene ring or amino group in different substrates (47, 52-56). This was also consistent with the catalytic oxidation process of CmO. Collectively, the mutation of individual amino acid residue was a self-defense behavior of microorganisms against the selection pressure of antibiotics, and also reduced the potential selection pressure of other antibiotics. These conserved amino acid residues in CmO and itself would be useful in the indication and evaluation of the resistance risk and biotransformation of CAP/TAP in diverse environments.

A homologous model was artificially constructed to reveal the catalytic mechanism of CmO at the molecular structure level, and N180, A183 and A209 amino acid residues were optimized by the molecular dynamics method. After the quality evaluation, the optimal pose was employed to calculate molecular docking and analyze the Grid score (−53.37 kcal/mol). The modelling results indicated that CAP was bound inside CmO to form a hydrophobic structure with the amino acid residues (A344, Y380 and M474) and small molecule FAD. The hydrophobic interaction exerted a considerable Van der Waals’ force to CAP binding (Grid vdw=-52.32 kcal/mol). Meanwhile, CAP could form a 3.0-3.5 Å hydrogen bond with the residues G99 (at the position of O atom), N518 and FAD through the C-3’ position, and form a 3.0 Å hydrogen bond with the phenolic hydroxyl group of the Y380 through the N atom of the amide bond. These hydrogen bonds determined the binding mode of CAP, however, electrostatic force only made a limited contribution (Grid es = -1.05 kcal/mol) due to the large length of the hydrogen bond (Figure 4A). In general, CAP binds to the CmO pocket at the C-3’ position relying on a well-defined binding force (Grid score = -53.37 kcal/mol) from the hydrogen bonds, which makes CAP preferentially catalyzed by preventing other substrates from binding.

**Figure 4.**
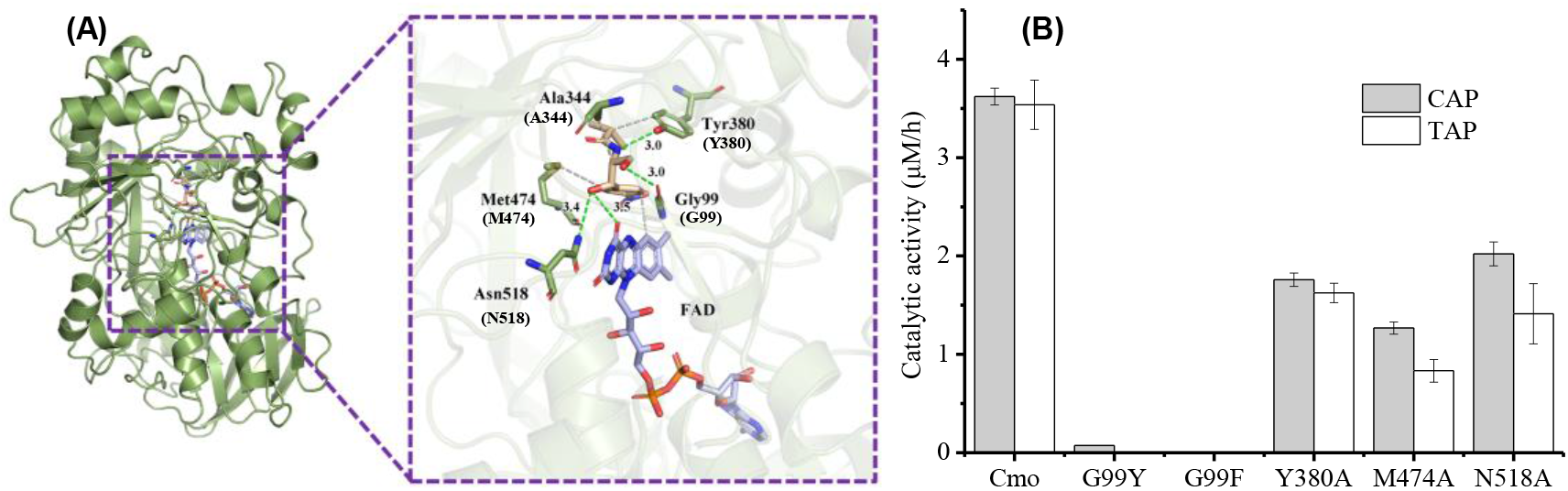
The 3D structure of CmO binding with CAP and activity analysis of CmO with site-directed mutation. The binding pattern of CAP and CmO protein: the smudge bands represent CmO protein; smudge, wheat, and light blue rods represent the key residues, CAP, and the coenzyme FAD, respectively; the green and gray dotted line indicate hydrogen bond and hydrophobicity, respectively (A). The comparison of catalytic activity of CmO at various amino acid residue mutation sites with CAP and TAP as the substrate (B).

The molecular docking results were verified by performing a site-directed mutagenesis of the key amino acid residues associated with the hydrogen bond forming. CmO after replacing the G99 by the phenylalanine (F) has lost complete oxidizing capacity of CAP and TAP (Figure 4B). The G99 was mutated to a large amino acid residue, which might break the bond between the O atom and the ligand, and in turn, being pushed out of the CmO pocket. Similarly, the G99Y variant reached only 2.01±0.28% of the original CAP catalytic activity. Furthermore, both the M474 and N518 residues were both mutated to the A residue, making the corresponding catalytic efficiencies be markedly decreased to 34.99-55.81% (CAP) and 39.94-45.93% (TAP) within 24 h. Due to the smaller volume of M474 and N518A variants, the binding between the enzyme and the ligand was weakened and the ligand binding pocket was deformed. The catalytic efficiencies of Y380A variant also decreased by 45.93-48.57% (Figure 4B) due to the replacement of the phenolic hydroxyl group initially bounding to the N atom of the CAP amide bond. Taken together, the mutations induced by the bulky amino acid at the Gly99 residue site could rigidly disrupt the binding between CAP/TAP and CmO, destroying the catalytic activity; moreover, the Y380A, M474A, and N518A variants containing the single-site mutations could restrain the formation of hydrophobic interaction to a certain degree, affecting the activity in consequence. These observations strongly suggested that the hydrogen bond formation plays a crucial role in the catalysis of CAP/TAP by CmO, while also matching the multiple alignment analysis.

### Comparison of CAP susceptibility in the *cmO* and other CAP-degrading resistance genes expressed in *E. coli*

Three target genes (an amide bond-hydrolyzing esterase gene *estDL136*(28), a nitroreductase gene *nfsB*(13), and a acetyltransferase gene *cat*(43)) were cloned into the expression vector pET29a(+) and expressed in *E. coli* to compare CAP susceptibility with *cmO*. The degradation efficiency of CAP was evaluated, and its intermediates were characterized by analyzing the transformants carrying the recombinant plasmids (Figure S9). The modified microbroth dilution susceptibility testing was operated to calculate 50% inhibitory concentration (IC_50_). The *E. coli* carrying an empty vector (negative control) was susceptible to CAP, with a calculated IC_50_ of 2.39±0.26 μg/mL and an observed MIC of 8 μg/mL (Figure 5AB). *E. coli* expressing *cmO* was highly resistant to CAP at an IC_50_ value of 18.59 ± 0.25 μg/mL and an observed MIC value of 32 μg/mL. Its susceptibility was second only to BL29cat with an IC_50_ at 25.57±2.07 μg/mL and a MIC at 64 μg/mL. Although the IC_50_ of *cat* in this experiment was lower than that in a previous study (over 256 μg/mL) (13), the susceptibility of *cmO* expression was second only to that of *cat* expression. The IC_50_ of other two strains with *nfsB* and *estDL136* genes expression were 9.14±0.23 and 4.16±0.06 μg/mL respectively, and the MIC of them were 16 and 8 μg/mL respectively. It was suggested that the order from the strong sequence to the weak was *cat, cmo, nfsB*, and *estDL136* by comparing the resistance variation of these four CAP-degrading resistance genes.

**Figure 5.**
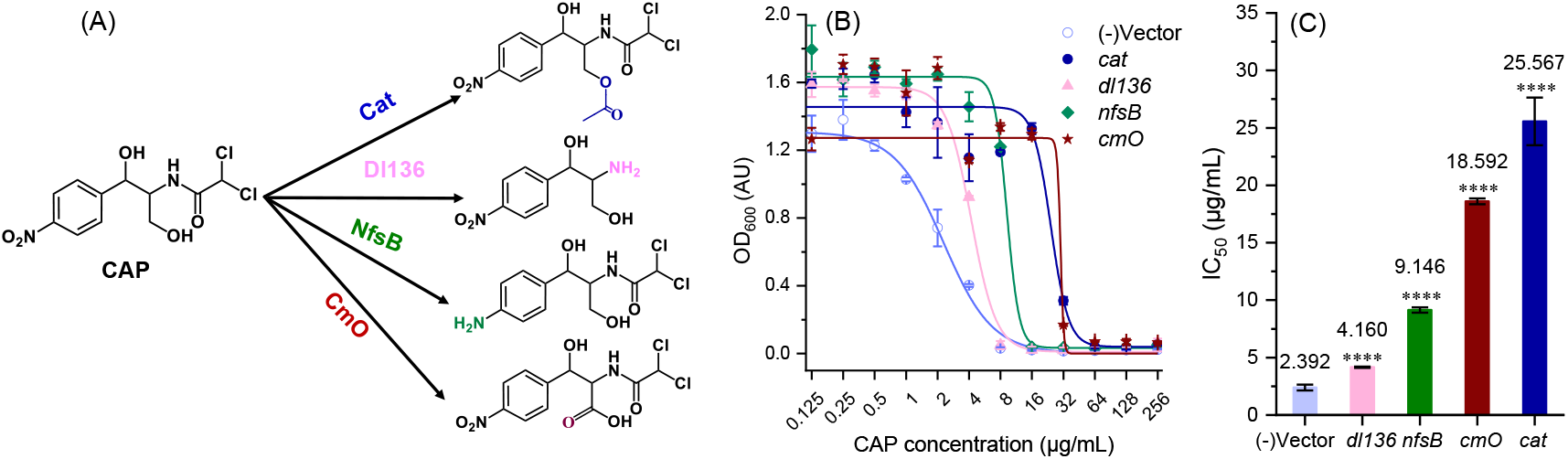
Four CAP resistance mechanisms mediated by enzymatic inactivation (A). The *cmO* and other CAP-degrading resistance genes expression can alter susceptibility of *E. coli* to CAP. Dose-response curves of microbroth dilution assays for *E. coli* expressing CAP-degrading resistance genes in the presence of CAP (A) with corresponding IC50 calculated from the curve fit (B). The “*(-)vector*” represents the empty vector of pET29a(+), “*cat*” represents the acetyltransferase gene, “*dl136*” represents the esterase gene, “*nfsB*” represents the nitroreductase gene in previous studies and “*cmO*” represents CAP oxidase gene in this study. All points are averages of 6 biological replicates experiments with standard deviation (SD) error bars. Statistical significance was calculated with respect to the vector control by the two-tailed unpaired t-test. Adjusted *P* value is displayed as ****p ≤ 0.0001

## MATERIALS AND METHODS

### Chemicals

CAP and TAP (both purity >99%) were purchased from Macklin (Shanghai, China). Other antibiotics and chemical reagents used were of the highest analytical purity.

### Bacterial strains, plasmids, primers and culture conditions

The bacterial strains, plasmids and the primers used in this study are listed in Table S1. *Sphingobium* sp. CAP-1 was grown at 30°C in Luria–Bertani (LB) medium or mineral salt medium (MSM, pH 7.0), each supplemented with CAP or TAP(16). *Escherichia coli* strains were grown at 37°C in LB medium, in which, antibiotics ampicillin (Amp, 100 mg/L), gentamicin (Gm, 100 mg/L), spectinomycin (Sm, 100 mg/L), streptomycin (Str, 100 mg/L) and kanamycin (Km, 50 mg/L) were added as required.

### Screening out a novel CAP oxidase in strain CAP-1

The comparative proteomic analysis of strain CAP-1 was previously carried out through isobaric tags for relative and absolute quantitation (iTRAQ) technology (16). Based on the genomic annotation and the fold change (FC) of the upregulated-expression proteins, one candidate gene was screened out (FC > 1.5 based, *P* < 0.05 on Student’s t-test, and the associated false discovery rate within 0.05 based on the Benjamin algorithm). The gene was further amplified with FastPfu DNA polymerase from genomic DNA of strain CAP-1 with the primer pair of ChoF/R (Table S1). The PCR products were ligated into pMD18-T vector and transformed into *E. coli* DH5α (32). The positive transformants were applied for the biotransformation assay. The oxidative biotransformation of CAP or TAP to O-CAP or O-TAP was confirmed using HPLC and high-resolution mass spectrometry as described below. A multiple alignment of the amino acid sequence of CmO and other biochemically identified oxidases (dehydrogenases) was performed as previously described (33, 34). Phylogenetic analysis of CmO and homologous oxidases was performed by using MEGA (version 7.0).

### Overexpression and purification of CmO

The *cmO* gene from the genomic DNA of strain CAP-1 was amplified with primers *cmO*-F/*cmO*-R (Table S1). The *cmO* fragment was ligated into a NdeI/XhoI-digested pET29a(+) plasmid using the ClonExpress II one-step cloning kit (Vazyme Biotech, China) to produce pET29a-*cmO*. The recombinant plasmid was subsequently transformed into *E. coli* BL21(DE3) and the positive transformant was cultured in LB medium containing Km (50 mg/L) at 37°C until the OD_600_ reached 0.6. Protein expression was induced at 16°C with 0.8 mM IPTG for 12 h. The cells of *E. coli* BL21 harboring pET29a-*cmO* were collected after centrifuging at 15,000g and 4°C for 20 min and washing twice with phosphate buffered saline (PBS, pH 7.4). The suspended cells were sonicated (Ultrasonic Cell Crusher, Thermo Fisher Scientific) and centrifuged under the conditions above to remove the cell debris. CmO was purified by using a Ni-nitrilotriacetic acid (Ni-NTA)-Sefinose column (Sangon Biotech, Shanghai, China) and dialyzed in Tris-HCl buffer (pH 7.4) overnight to remove imidazole and Ni^2+^. The purity and molecular mass of CmO were shown by SDS-PAGE, and the protein concentration was determined with the UV absorption peak at 280 nm by Nanodrop One (Thermo Scientific, USA).

### Enzyme assay

The enzyme activity was measured in 1 mL PBS (10 mM, pH=7.4) supplemented with the proper amount of purified CmO at 30°C for 10-20 min in triplicate. CAP and TAP were used as the reaction substrates with various initial concentrations, separately. Each enzymatic reaction was terminated by boiling for 1 min, and diluted by the equal volume methanol. Controls with inactive enzyme were also carried out under the same conditions. The determination of kinetic parameters including *V*_max_ and *K*_m_ were described elsewhere (35).

To determine the optimum temperature of CmO, the relative activities were compared at 10-80°C. In different buffers (36), the optimum pH was determined by comparing the enzyme activities in the pH range from 3.8 to 10.6. To determine the stability, CmO was pretreated in different temperature and pH for 1 h to analyze the remaining activity, respectively. The effects of metal ions, inhibitors and detergents on the enzyme activity were analyzed as previously described(35). The concentration of all metal ions and inhibitors were 1 mM. The detergents that comprised Triton X-100, Tween-20, Tween-80, sodium dodecyl sulfate (SDS) and cetyltriethyl ammonium bromide (CTAB) were added at a final concentration of 0.1% (m/V).

### Homology modeling and molecular docking

The 3D structure of the target protein, CmO (UniProtKB AC: A0A6I5YGT3), was constructed using an online protein structure prediction server, SWISS-MODEL (http://swissmodel.expasy.org/). The template protein was identified through BLAST sequence search. The molecular dynamics (MD) simulations were performed in Yinfo Cloud Computing Platform (YCCP, https://cloud.yinfotek.com/) to optimize the protein structure. The quality of protein model was evaluated using the online website SAVES V6 (https://saves.mbi.ucla.edu/) (35). Molecular docking was conducted in Yinfo Cloud Computing Platform. By the DOCK 6.9 program, CAP was docked into the enzyme CmO pocket. Based on the quality of the protein, the optimum binding mode was selected for analysis (37-39).

### Site-directed mutagenesis

The mutations were introduced into CmO (expressed on pET-29a(+)) using a Mut Express II Fast Mutagenesis Kit (Vazyme, Nanjing, China) according to the manufacturer’s protocols. The primers used for site-directed mutagenesis of CmO were shown in Table S1, and the mutant amino acid residues included glycine (G)99, tyrosine (Y)380, methionine (M)474 and asparagine (N)518.

### Expression of CmO in *Sphingomonas wittichii* RW1

The fragment of *cmO* was amplified from the genomic DNA of strain CAP-1 with the primers pBBcmoF/R (Table S1). The PCR product was then ligated into KpnI/EcoRI-digested broad-host-range plasmid pBBR1-MCS2 using the ClonExpress II one-step cloning kit (Vazyme Biotech, China) to generate pBBR-*cmO*, which was electroporated into *S. wittichii* RW1 (a non-CAP degrader), and selected on LB plates supplemented with Sm (100 mg/L) and Km (50 mg/L).

### Analytical methods

CAP, TAP, O-CAP and O-TAP were analyzed using a HPLC (model-1260, Agilent Technologies, USA) equipped with a C_18_ column (4.6×100 mm, 2.7 μm, Agilent Technologies, USA). The mobile phase was a mixture of acetonitrile and ultrapure water containing 0.1% formic acid (30:70; vol/vol) at a flow rate of 0.6 mL/min in isocratic mode. A UV detector (model-1260, Agilent Technologies, USA) was set at 245 nm and 275 nm. For the refined identification of the O-TAP and O-CAP chemical structure, UPLC-triple-quadrupole mass spectrometer (UPLC-TQMS) was operated with appropriate parameters mentioned elsewhere(40). The high-resolution mass spectrometry and nuclear magnetic resonance (NMR) analysis using a 700 MHz liquid NMR spectrometer (AVANCE III, Bruker Biospin, Switzerland) were employed as described previously (16). The sample for NMR was separated by a high performance preparative liquid chromatography, freeze-dried to a solid powder and locked with deuterated chloroform (CDCl_3_) (16, 41, 42).

### Calculation of IC_50_ toward to CAP with different CAP resistant strains

Three types of CAP resistance via enzymatic inactivation including an amide bond-hydrolyzing esterase gene *estDL136* (28), a nitroreductase gene *nfsB* (13), and a acetyltransferase gene *cat* (43), were commercially synthesized and cloned into the multiple clone sites of *Nde*I/*Xho*I in vector pET-29a(+) (GenScript, USA), and then were transformed into *E. coli* BL21(DE3) to generate BL29dl136, BL29nfsB, and BL29cat, respectively. Microbroth serial dilution assays were employed to determine IC_50_ (half maximal inhibitory concentration) values for CAP and in a modification of EUCAST standard protocol. 100 μL volumes of Mueller-Hinton broth (MHB) were added into 96-well plates with CAP concentrations ranging from 0.125 to 256 μg/mL. The above constructed strains and BL29cmO were cultured overnight in MHB with Km and inoculated into fresh MHB to the OD_600_ value of 0.1. Diluted cultures were added to the prepared CAP-containing 96-well plates in 100 μL aliquots for an initial OD_600_ and volume of 0.05 and 200 μL, respectively. Growth with 6 biological replicates in the microplate constant temperature shaker at 800 rpm for 36 h at 30°C was measured by a Spark 10M multimode microplate reader (Tecan, Switzerland). Dose-response curves and significant differences in IC_50_ were analyzed and calculated as described elsewhere (13).

## Data availability statement

The gene *cmO* and genome sequence of *Sphingobium* sp. CAP-1 and *Sphingopyxis* sp. GC21 have been deposited in the GenBank database under accession numbers of OP019282, OP019283, CP046252−CP046256 and CP095845, respectively.

## Supplemental Material

Supplemental material is available online only. Detailed description of the strains, plasmids, primers, six NMR spectra (^1^H, ^13^C, DEPT, HMQC, HMBC, and ^1^H−^1^H COSY), the HR-ESIMS spectrum of O-TAP, the CAP/TAP degradation characteristics of recombinant *S. wittichii* strain RW1-*cmO* and strain BL29cmo, the SDS-PAGE picture and enzymatic catalytic characteristics of purified CmO, the amino acid sequence alignment of CmO with several oxidases belonging to the GMC family and the biotransformation characteristics of the transformants carrying three plasmids (pET29a-*cat*, pET29a-*estDL136* and pET29a-*nfsB*).

## ACKNOWLEDGMENTS

The study was financially supported by the National Natural Science Foundation of China (Nos. 31870102, 52170188 and 52091545), Shenzhen Overseas High-level Talents Research Startup Program (No. 20210308346C), and Shenzhen Science and Technology Program (No. KQTD20190929172630447).

