## Supplemental Table 1, Supplemental Figure 1-9 for "The Molecular Mechanism of Chloramphenicol and Thiamphenicol Resistance Mediated by a Novel Oxidase CmO in Sphingomonadaceae"

**TABLE S1** The strains, plasmids and the primers used in this study

| Strain | Relevant characteristic(s) ^a^ | Source or reference |
| --- | --- | --- |
| *Sphingobium* sp*.* CAP-1 | Wild type; Str^r^; capable of catabolizing CAP/O-CAP/PNBD/PNBA | (1) |
| *Sphingomonas* sp. RW1 | Dibenzo-p-dioxin-degrading stain; Sm^r^ | (2) |
| *Sphingopyxis* sp. GS21 | Wild type; Km^r^, Gm^r^, Str^r^; capable of catabolizing CAP/O-CAP/PNBD/PNBA | This study |
| *E. coli* DH5α | *supE44λ- thi-1 gyrA96 relA1 phoA;* Φ80d/*lac*ZΔM15 *Δ(lacZYA-argF)*U169*;* *endA1 recA1hsdR17(rk-,mk+)* | Tiangen |
| *E. coli* BL21(DE3) | F- *ompT hsdSB(rB- mB-) gal dcm* (DE3) | Tiangen |
| *Sphingomonas wittichii* RW1*-cmO* | The mutant strain with pBBR-*cmO*; capable of catabolizing CAP/O-CAP/PNBD/PNBA | This study |
| Plasmid Relevant characteristic(s) ^a^ Source or reference | | |
| pMD18-T | Transformed from plasmid pUC18 vector; *Lacz* *Amp^r^, sacB* | TaKaRa |
| pM18-*cmO* | The cmo gene cloned in pMD18-T, Km^r^ | This study |
| pET-29a(+) | IPTG inducible expression vector, Km^r^ | Novagen |
| pET29-*cmO* | The cmo gene cloned in pET-29a(+), Km^r^ | This study |
| pBBR1-MCS2 | Broad-host-range cloning vector, Km^r^ | (3) |
| pBBR-*cmO* | The *cmo* gene cloned in pBBR1-MCS2, Km^r^ | This study |
| pET29a-*cat* | The *cat* gene cloned in pET-29a(+), Km^r^ | This study |
| pET29a- *estDL136* | The *estDL136* gene cloned in pET-29a(+), Km^r^ | This study |
| pET29a- *nfsB* | The *nfsB* gene cloned in pET-29a(+), Km^r^ | This study |

| Function | Primers | Restriction sites | Sequence (5’ to 3’) |
| --- | --- | --- | --- |
| Cloning and verification | ChoF | None | tcacatggagggcattgtgcaaga  tcagtggctttttcggatcagatc |
|  | ChoR | None |  |
| Plasmid recombination for expression | *cmO*-F | NdeI | gtgccgcgcggcagccatatggagggcattgtgcaagatattaga |
|  | *cmO*-R | XhoI | gtggtggtggtggtgctcgaggtggctttttcggatcagatcg |
| Plasmid recombination for CmO site-directed mutation | G99F-F | None | gctcgtcgatcaactttatgatgtatgttcgcggcaat |
|  | G99F-R | None | aaagttgatcgacgagccgccgccgaggactt |
|  | G99Y-F | None | aaagttgatcgacgagccgccgccgaggactt |
|  | G99Y-R | None | atagttgatcgacgagccgccgccgaggactt |
|  | Y380A-F | None | tatacctggcgaagcgtgcagcaattggcgtt |
|  | Y380A-R | None | acgcttcgccaggtatacgccgtccggcgtga |
|  | M474A-F | None | cttcctggcgtatcacccgactggcacttgcg |
|  | M474A-R | None | ggtgatacgccaggaaggcgctctggcggata |
|  | N518A-F | None | tcagcgcggcgacaaatgcaccgtgcatcatg |
|  | N518A-R | None | atttgtcgccgcgctgaccagcgtcggcatga |
| Plasmid recombination for gene complementation | pBBcmoF | KpnI | aaagggaacaaaagctgggtaccatggagggcattgtgcaaga |
|  | pBBcmoR | EcoRI | atcccccgggctgcaggaattctcagtggctttttcggatcag |

^a^ Str^r^, Sm^r^, Km^r^, Gm^r^ and Amp^r^ resistance to streptomycin, spectinomycin, kanamycin, gentamicin and ampicillin, respectively.

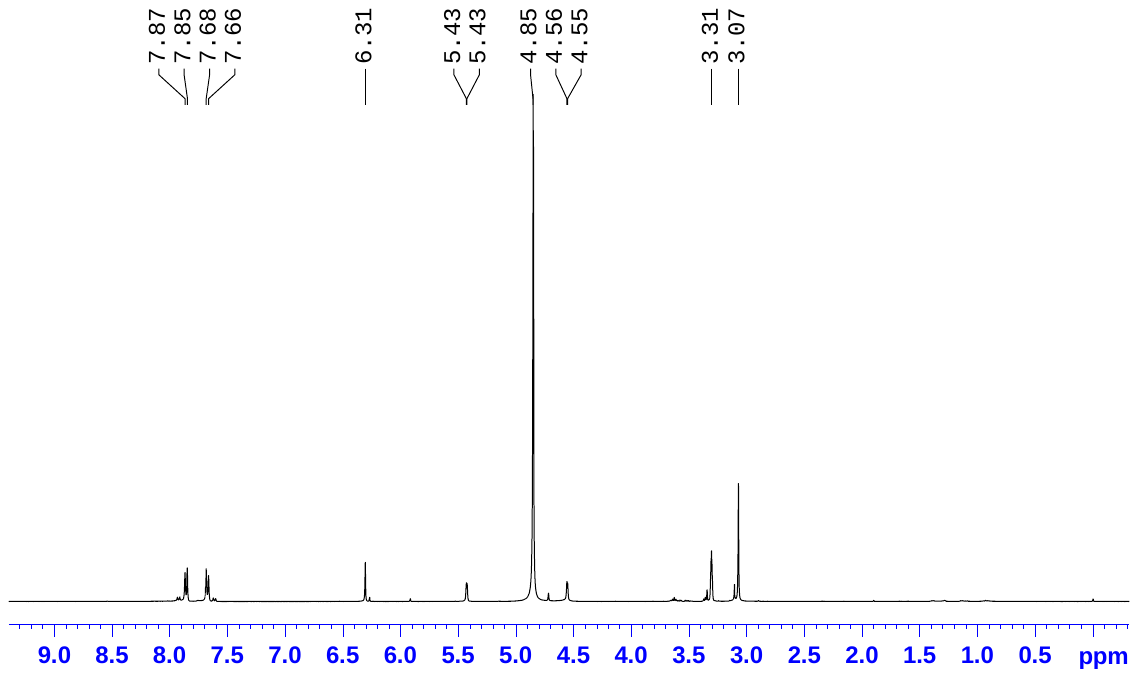

**Fig. S1A** ^1^H NMR spectrum for compound O-TAP

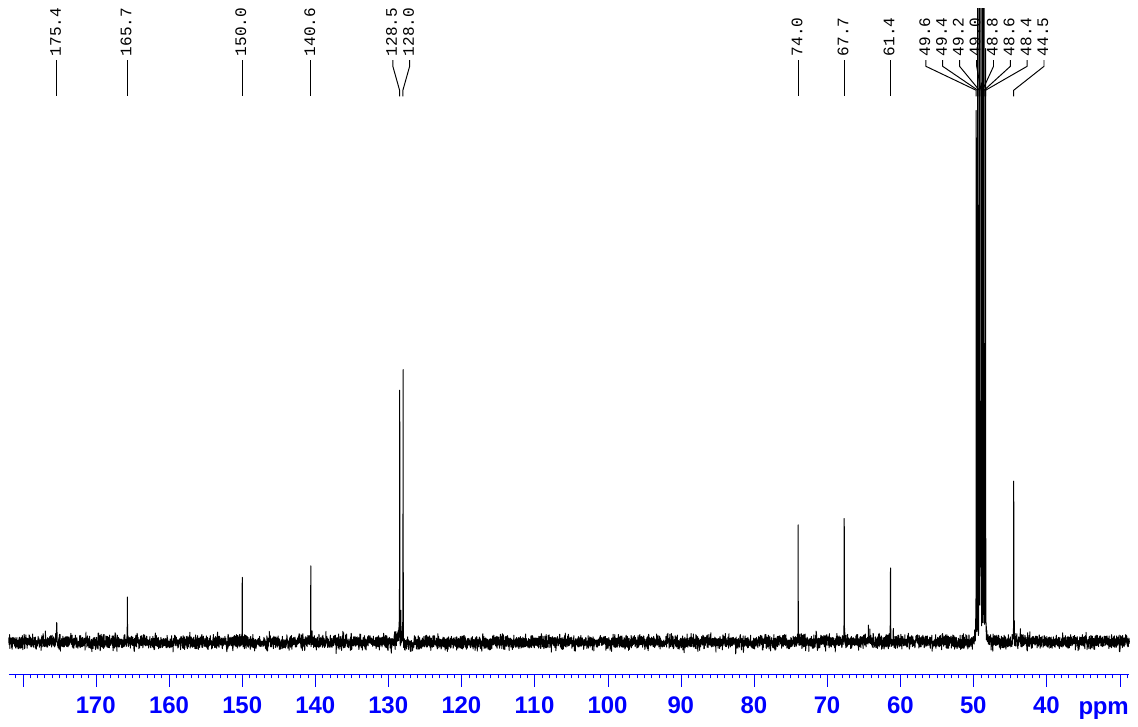

**Fig. S1B** ^13^C NMR spectrum for compound O-TAP

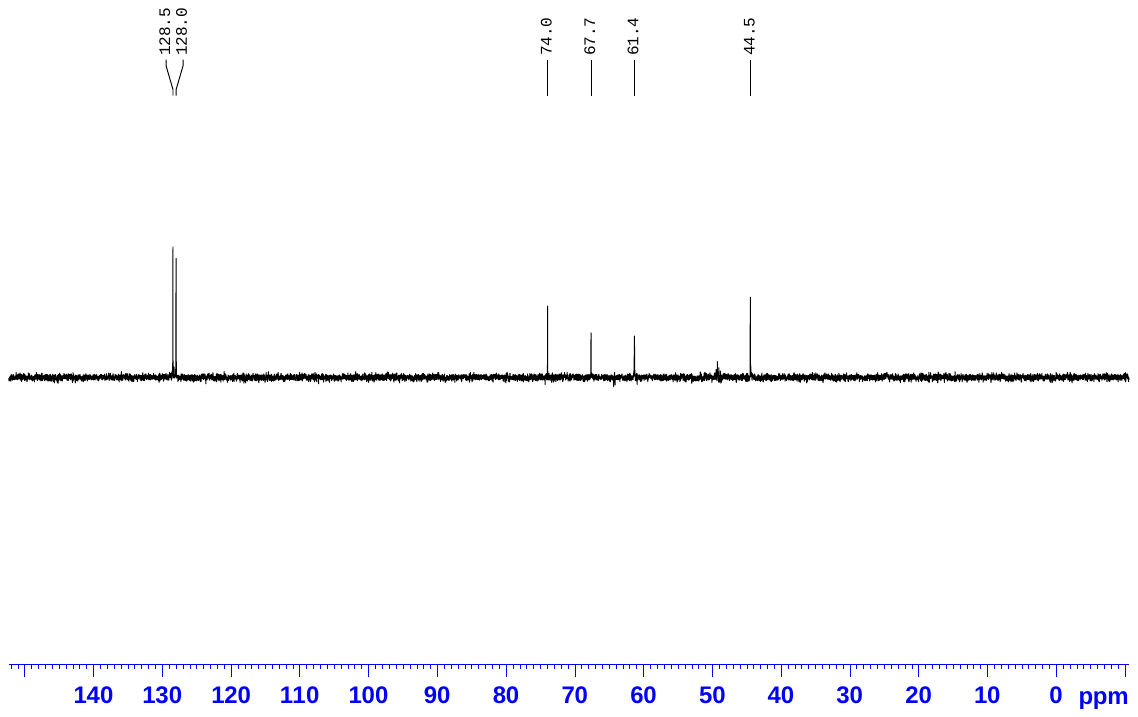

**Fig. S1C** DEPT (*θ=135˚*) NMR spectrum for compound O-TAP

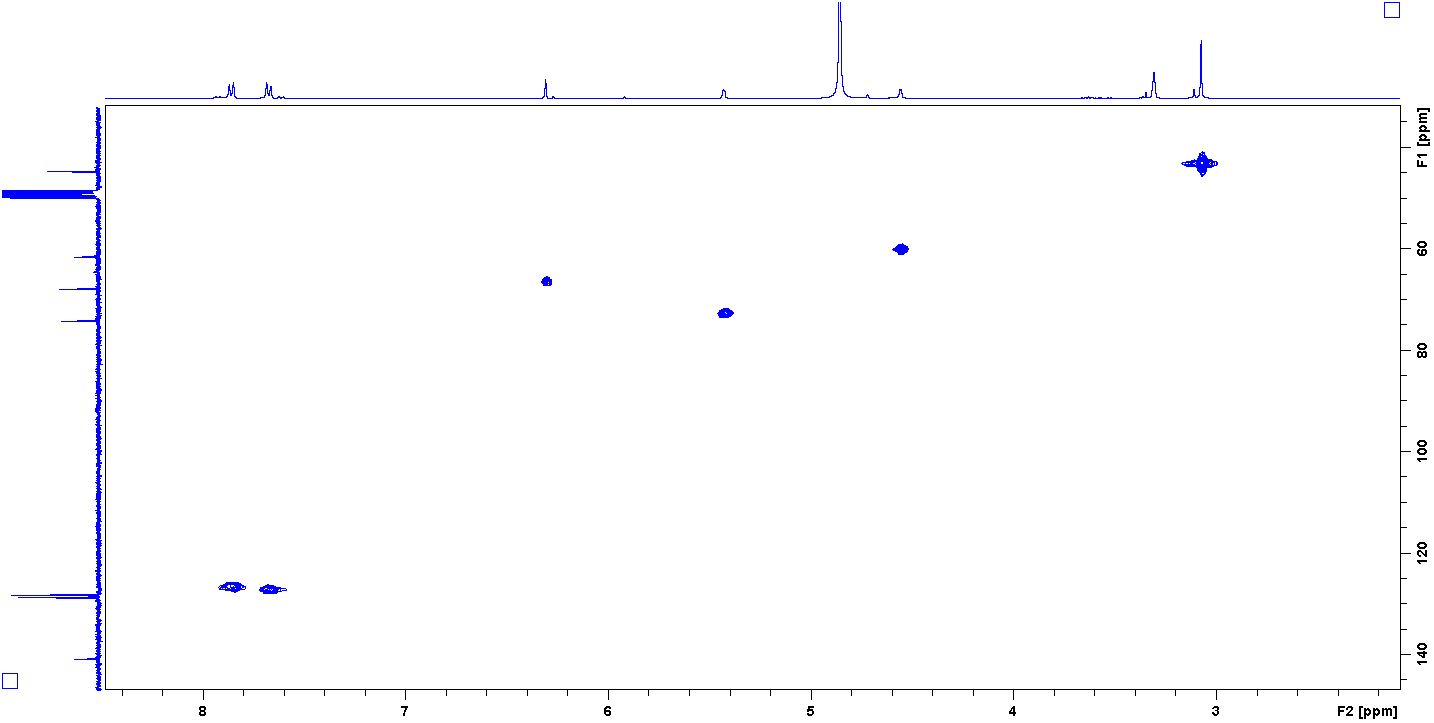

**Fig. S1D** HMQC spectrum for compound O-TAP

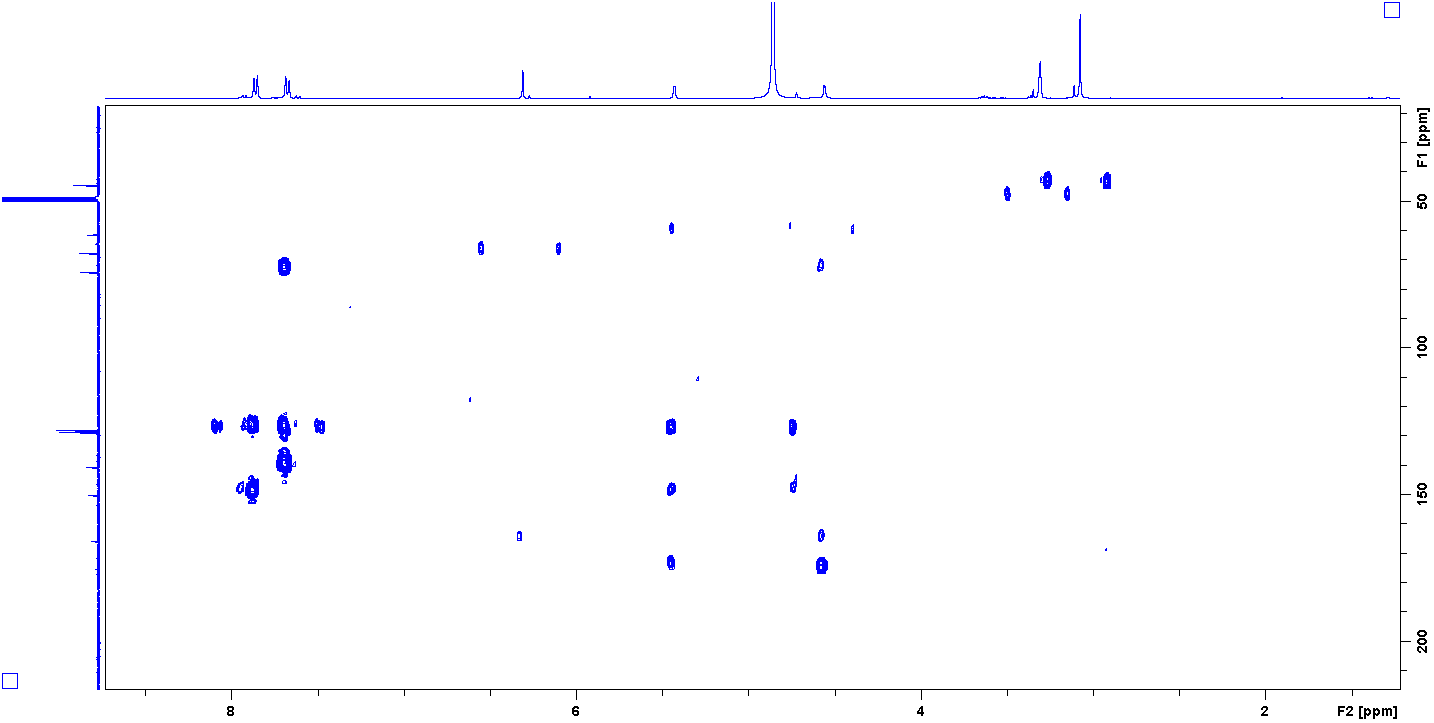

**Fig. S1E** HMBC spectrum for compound O-TAP

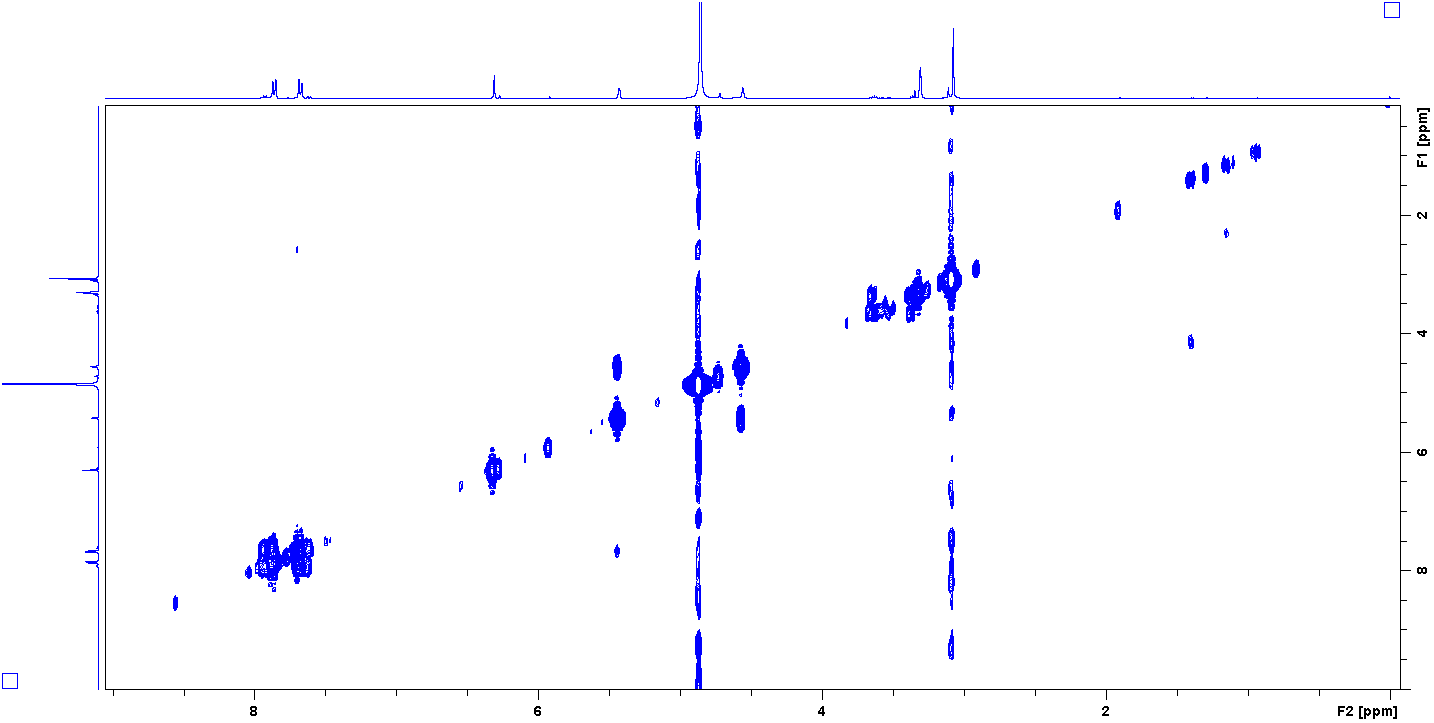

**Fig. S1F** ^1^H**−**^1^H COSY spectrum for compound O-TAP

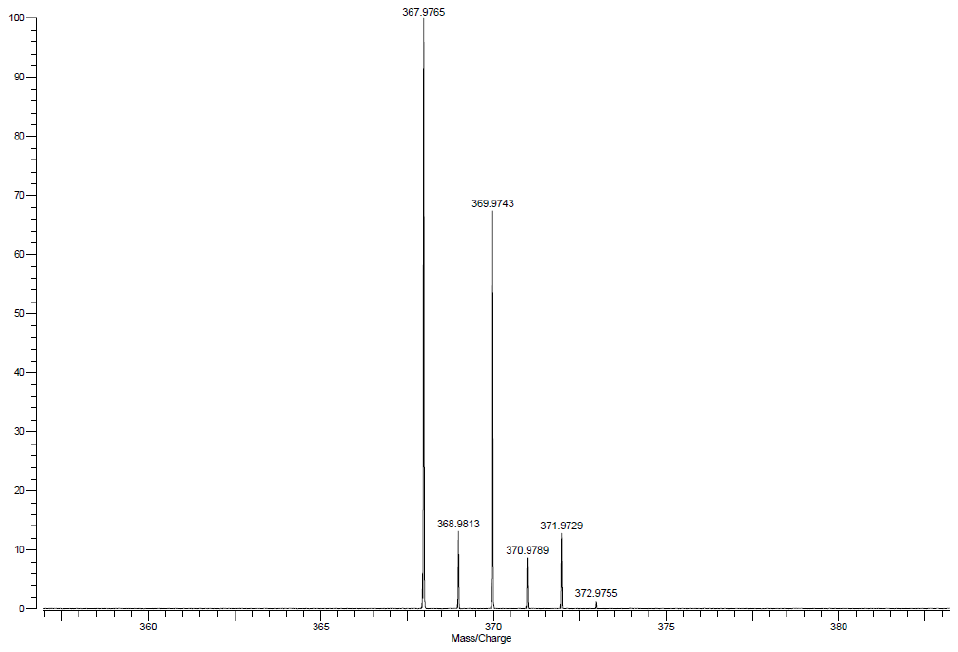

**Fig. S1G** HR-ESIMS spectrum for compound O-TAP

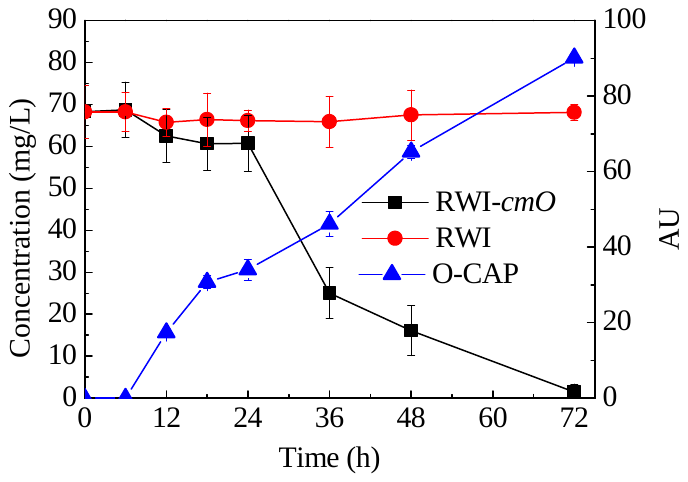

**Fig. S2** Comparation of CAP degradation characteristics between *S. wittichii* RW1 and the recombinant strain RW1-*cmO*.

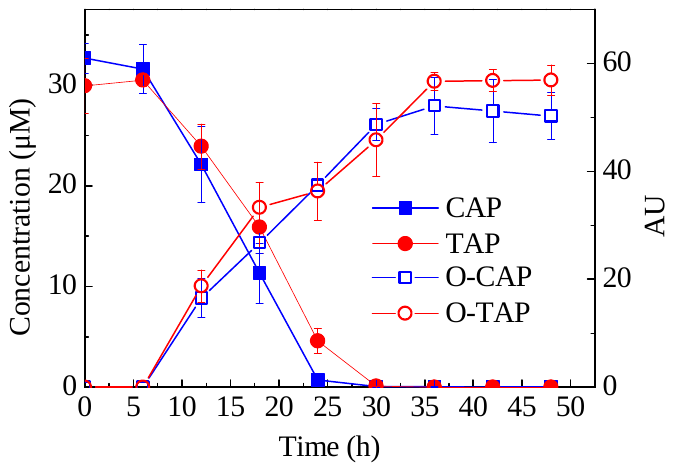

**Fig. S3** Degradation characteristics of CAP and TAP by strain BL29cmo.

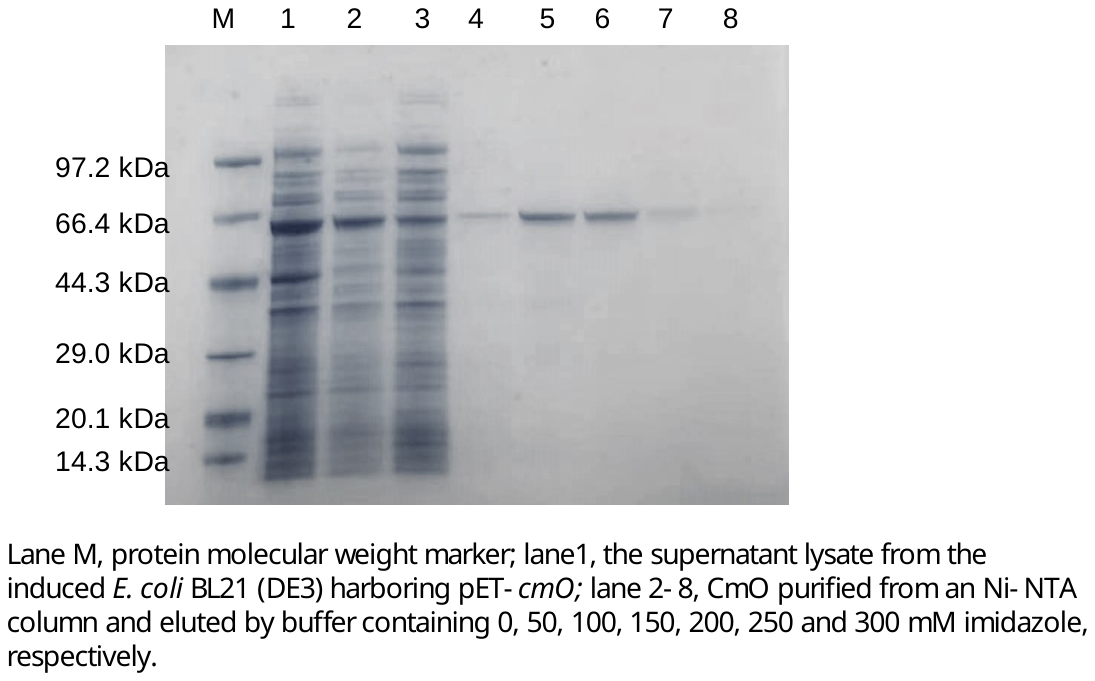

**Fig. S4** Lane M, protein molecular weight marker; lane1, the crude extract from the induced *E. coli* BL21 (DE3) harboring pET29a-*cmO*; lane 2-8, CmO purified from an Ni-NTA column and eluted by buffer containing 0, 50, 100, 150, 200, 250 and 300 mM imidazole, respectively.

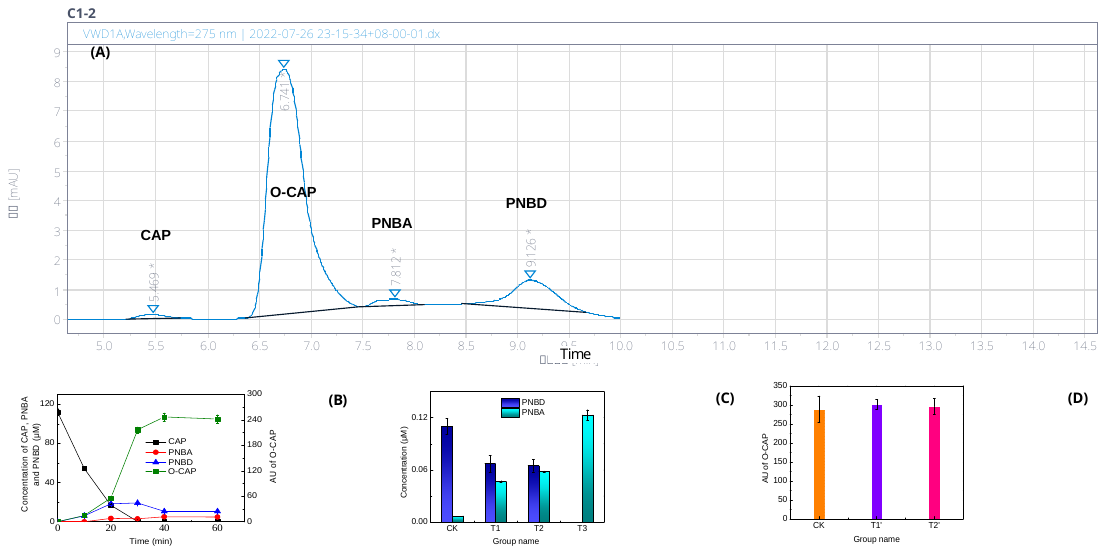

**Fig. S5** The HPLC chromatogram of 100 μM CAP degradation products (30 min): O-CAP, PNBD and PNBA (A), the biotransformation characteristics of CAP (B), PNBD (C) and O-CAP (D) catalyzed by the purified CmO. CK represents the reaction system with inactivated CmO; T1 and T1’ represent the reaction system with 73.89 μg/L CmO; T2 and T2’ represent the reaction system with 147.78 μg/L CmO; T3 represents the reaction system with sufficient crude extract (150 μL).

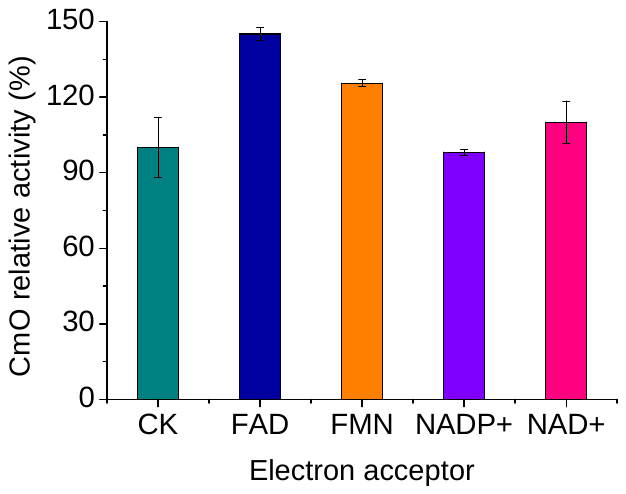

**Fig. S6** Effects of different electron acceptors on CmO relative activity.

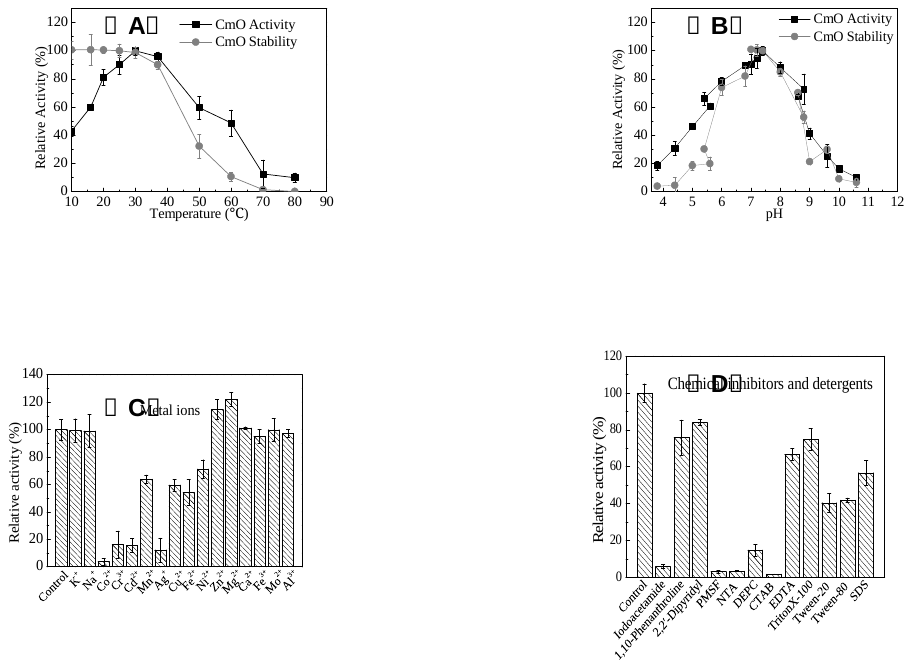

**Fig. S7** Effects of temperature (A), pH (B), different metal ions (C) and chemical inhibitor/detergents (D) on the catalytic activity and stability of CmO.

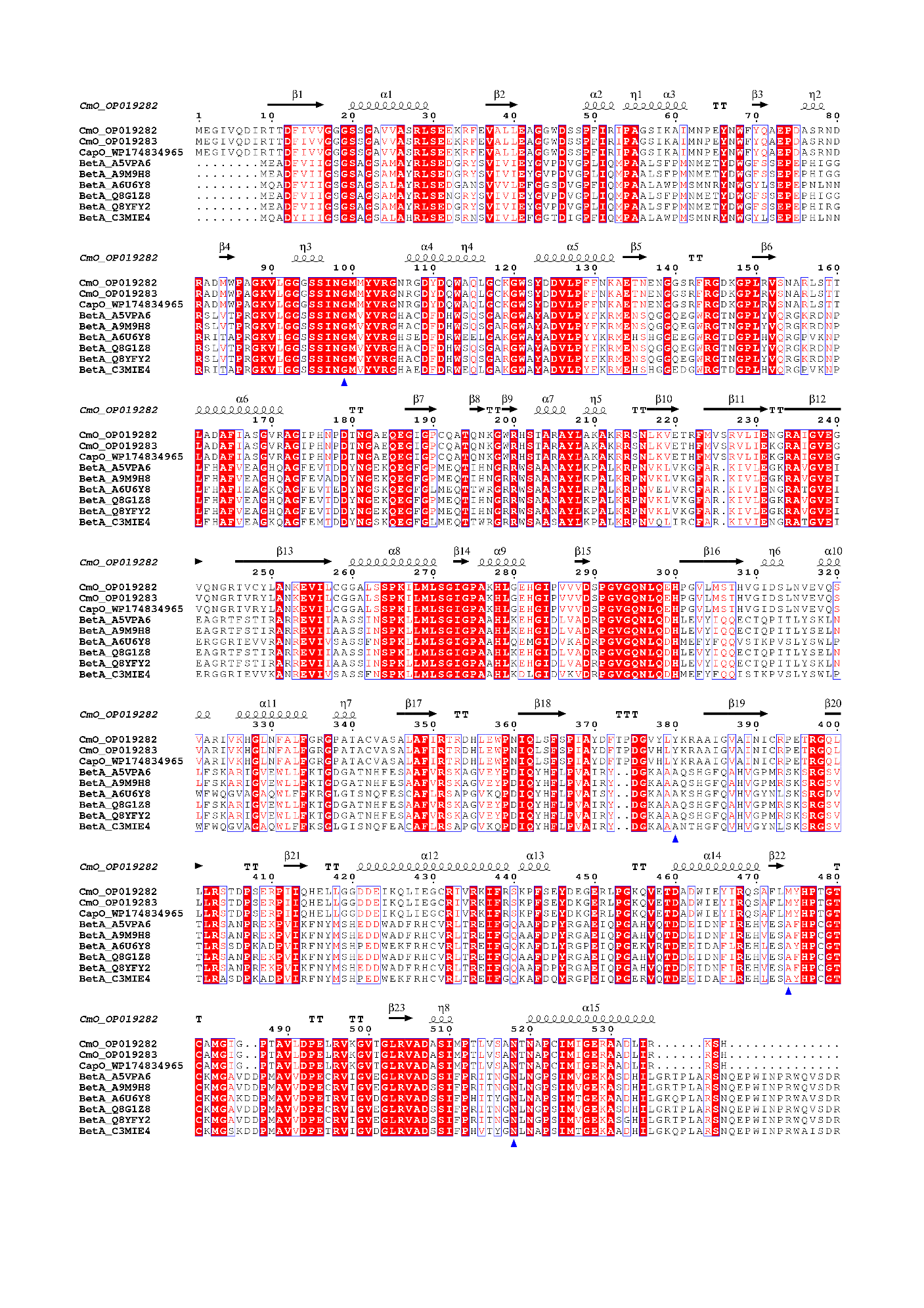

**Fig. S8** Amino acid sequence alignment of CmO with several oxidases belonging to the GMC family (cover 97%; 40.87 to 41.40% identities). The oxidases from *Sphingopyxis* sp. GS21 and *Sphingomonas* sp. CL5.1 were also aligned with CmO in *Sphingobium* sp. CAP-1 with the high amino acid homology (> 98.8%). Identical amino acids are displayed in white font on a red. The amino acid residue sites of hydrogen bond formation, using for site-directed mutagenesis analysis in Figure 4, were noted by blue triangle.

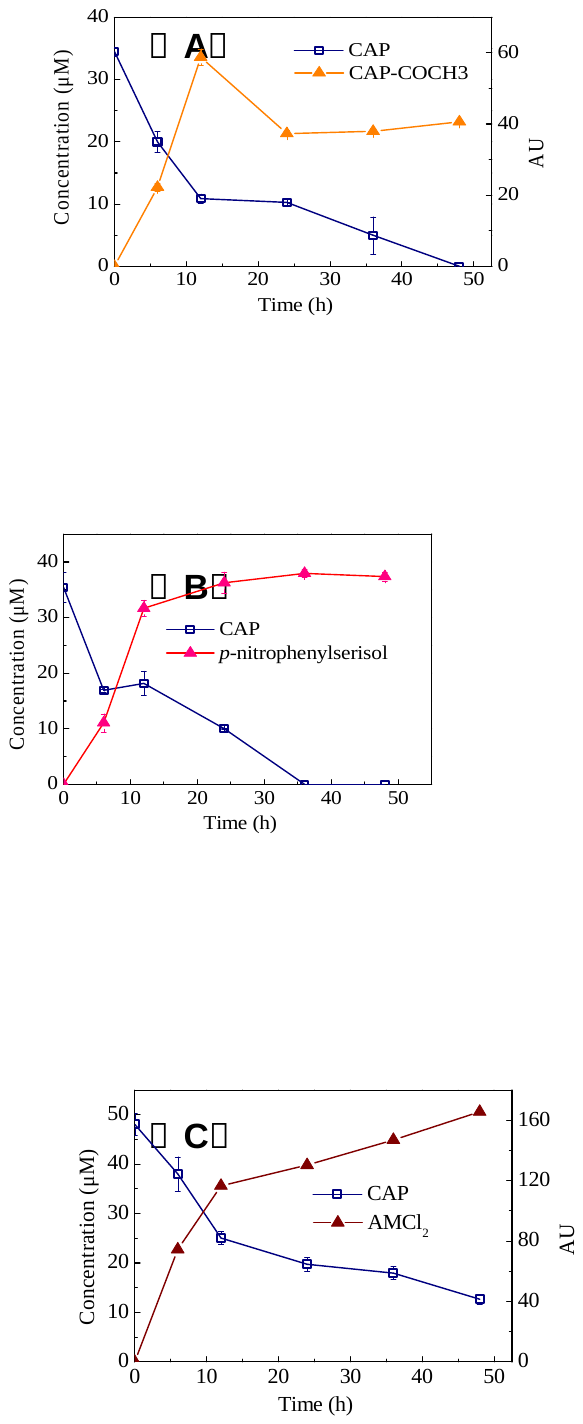

**Fig. S9** Degradation characteristics of the transformants carrying three plasmids: pET29a-*cat*(A), pET29a-*estDL136* (B) and pET29a- *nfsB* (C).

**OP019282** (The oxidase gene *cmO* to catalyze CAP and thiamphenicol in *Sphingobium* sp. CAP-1)

TCAGTGGCTTTTTCGGATCAGATCGGCTGCCCGTTCGCCAATCATGATGCACGGTGCATTTGTATTCGCGCTGACCAGCGTCGGCATGATCGAGGCATCCGCGACGCGAAGACCGGTGACGCCCTTGACGCGCAACTCCGGATCGAGGACCGCGGTCGGCCCAATTCCCATCGCGCAAGTGCCAGTCGGGTGATACATCAGGAAGGCGCTCTGGCGGATATACTCGATCCAATCAGCGTCGGTTTCGACCTGCTTTCCGGGCAGGCGTTCACCTTCGTCATATTCACTGAATGGCTTGGAACGGAAAATCTTGCGCACGATCCGACATCCTTCGATGAGCTGCTTGATCTCATCATCTCCTCCGAGCAGCTCATGTTGGATAATCGGCCGCTCGCTTGGATCGGTGGAGCGGAGCAGTAGCTGACCGCGCGTCTCGGGCCGGCAGATGTTGATGGCAACGCCAATTGCTGCACGCTTGTACAGGTATACGCCGTCCGGCGTGAAGTCGTACGCGATCGGCGAGAACGACAGTTGGATGTTGGGCCACTCGAGATGGTCCCGCGTGCGAATGAACGCGAGAGCGGAGGCAACACATGCCGTGGCTGGCCCTCGCCCGAACAAAGCGAAGTTCAAGCCATGCTTGACTATCCTGGCGACGCTTTGCACTTCGACATTGAGGCTATCGATGCCGACATGGGTCGACATCAACACTCCCGGATGTTCCTGCAGATTTTGCCCCACTCCCGGGGAATCGACGACAACTGGGATGCCATGCTCGCCAAGATGCTTTGCCGGGCCAATGCCCGAGAGCATCAATATCTTCGGCGACGACAAGGCGCCGCCGCAAAGAATGACCTCCTTGTTTGCCAGGTAGCAAACCGTGCGCCCGTTCTGAACGCCTTCGACGCCGATCGCGCGGCCGTTCTCGATCAGTACCCGACTGACCATGAAACGCGTCTCGACCTTCAAATTGGACCGGCGCTTCGCCTTGGCCAGATAGGCACGTGCCGTTGAATGTCGCCAACCCTTGTTCTGGGTGGCTTGGCAGGGGCCGATACCCTCTTGCTCGGCACCGTTGGTATCCGGATTGTGCGGAATCCCCGCACGTACGCCAGAAGCGATGAATGCGTCGGCCAACGTGGTCGATAGGCGGGCATTCGATACGCGCAGAGGGCCCTTGTCGCCGCGAAAGCGCGAGCCGCCGTTTTCGTTCGTCTCGGCCTTGTTGAAGAACGGAAGCACGTCGTCATAGGACCAGCCCTTGCAGCCGAGCTGGGCCCATTGATCATAATCCCCGCGATTGCCGCGAACATACATCATCCCGTTGATCGACGAGCCGCCGCCGAGGACTTTGCCGGCCGGCCACATGTCTGCTCGATCGTTTCGCGAGGCATCCGGTTCCGCTTGATAGAACCAGTTGTACTCGGGGTTCATGATCGCCTTGATCGAGCCCGCCGGAATCCGGATGAAAGGCGAGCTGTCCCACCCGCCTGCTTCGAGCAACGCGACCTCAAAACGCTTCTCTTCGCTCAGCCGAGATGCGACCACCGCGCCACTCGAACCGCCCCCAACGACGATAAAGTCCGTAGTTCTAATATCTTGCACAATGCCCTCCAT

MEGIVQDIRTTDFIVVGGGSSGAVVASRLSEEKRFEVALLEAGGWDSSPFIRIPAGSIKAIMNPEYNWFYQAEPDASRNDRADMWPAGKVLGGGSSINGMMYVRGNRGDYDQWAQLGCKGWSYDDVLPFFNKAETNENGGSRFRGDKGPLRVSNARLSTTLADAFIASGVRAGIPHNPDTNGAEQEGIGPCQATQNKGWRHSTARAYLAKAKRRSNLKVETRFMVSRVLIENGRAIGVEGVQNGRTVCYLANKEVILCGGALSSPKILMLSGIGPAKHLGEHGIPVVVDSPGVGQNLQEHPGVLMSTHVGIDSLNVEVQSVARIVKHGLNFALFGRGPATACVASALAFIRTRDHLEWPNIQLSFSPIAYDFTPDGVYLYKRAAIGVAINICRPETRGQLLLRSTDPSERPIIQHELLGGDDEIKQLIEGCRIVRKIFRSKPFSEYDEGERLPGKQVETDADWIEYIRQSAFLMYHPTGTCAMGIGPTAVLDPELRVKGVTGLRVADASIMPTLVSANTNAPCIMIGERAADLIRKSH

**OP019283** (The oxidase gene *cmO* to catalyze CAP and thiamphenicol in *Sphingopyxis* sp. GC21)

TCAGTGGCTTCTTCGGATCAGATCGGCCGCCCGTTCGCCAATCATGATGCACGGTGCATTTGTATTCGCGCTAACCAGCGTCGGCATGATCGAGGCATCCGCAACGCGAAGACCGGTGACGCCCTTGACGCGCAACTCCGGATCGAGAACCGCTGTCGGCCCAATTCCCATCGCGCAAGTGCCAGTCGGGTGGTACATCAGGAAGGCGCTCTGACGGATATACTCGATCCAATCAGCGTCGGTTTCGACCTGCTTTCCGGGTAAGCGTTCACCTTTGTCATATTCACTGAATGGCTTGGAACGGAAAATCTTGCGCACGATCCGGCATCCTTCGATGAGCTGCTTGATCTCATCATCTCCGCCGAGCAGCTCATGTTGGATAATCGGCCGCTCACTTGGATCGGTGGAGCGGAGCAGCAACTGACCGCGCGTCTCGGGCCGGCAGATGTTGATGGCAACGCCAATTGCCGCACGCTTGTACAGGTGTACGCCGTCCGGCGTGAAGTCGTACGCGATCGGCGAGAACGACAGTTGGATGTTGGGCCACTCGAGATGGTCTCGCGTGCGAATGAACGCGAGAGCGGAGGCAACGCATGCCGTGGCTGGCCCTCGCCCAAACAAAGCGAAGTTCAAGCCATGCTTGACTATCCTGGCGACGCTTTGCACTTCGACATTGAGGCTATCGATGCCGACATGGGTCGACATCAACACTCCGGGATGTTCCTGCAGATTTTGCCCCACTCCCGGGGAATCGACGACAACAGGGATGCCATGCTCGCCAAGATGCTTTGCCGGGCCAATGCCCGAGAGCATCAATATTTTCGGCGACGACAACGCGCCGCCGCAAAGAATGACCTCCTTGTTTGCCAAGTAGCGAACCGTGCGCCCGTTCTGAACGCCTTCGACGCCGATCGCGCGGCCTTTCTCGATCAGTACCCGACTGACCATGAAATGCGTCTCGACCTTCAGATTGGATCGGCGCTTCGCCTTGGCCAGATAGGCGCGTGCCGTTGAATGTCGCCAACCCTTGTTCTGGGTGGCTTGGCAGGGGCCGATACCCTCTTGCTCGGCACCGTTGGTATCCGGATTGTGCGGAATCCCCGCACGTACGCCAGAAGCGATGAATGCGTCGGCCAACGTGGTCGATAGGCGGGCATTCGATACGCGCAGAGGGCCCTTGTCGCCGCGAAAGCGCGAGCCGCCGTTTTCGTTCGTCTCGGCCTTGTTAAAGAACGGAAGCACGTCGTCATAGGACCAGCCCTTGCAGCCGAGCTGAGCCCATTGATCATAATCGCCGCGATTGCCGCGAACATACATCATCCCGTTGATCGACGAGCCGCCGCCGAGGACTTTGCCGGCCGGCCACATGTCTGCTCGATCGTTTCGTGAGGCATCCGGTTCCGCTTGATAGAACCAGTTGTACTCAGGATTCATGATCGCTTTGATCGAGCCCGCCGGAATCCGAATGAAAGGCGAGCTGTCCCACCCGCCGGCTTCGAGCAACGCGACCTCAAAACGCTTCTCTTCGCTTAGCCGAGATGCGACCACCGCGCCACTCGAACCGCCCCCAACGACGATAAAGTCCGTAGTTCTAATATCTTGCACAATGCCCTCCAT

MEGIVQDIRTTDFIVVGGGSSGAVVASRLSEEKRFEVALLEAGGWDSSPFIRIPAGSIKAIMNPEYNWFYQAEPDASRNDRADMWPAGKVLGGGSSINGMMYVRGNRGDYDQWAQLGCKGWSYDDVLPFFNKAETNENGGSRFRGDKGPLRVSNARLSTTLADAFIASGVRAGIPHNPDTNGAEQEGIGPCQATQNKGWRHSTARAYLAKAKRRSNLKVETHFMVSRVLIEKGRAIGVEGVQNGRTVRYLANKEVILCGGALSSPKILMLSGIGPAKHLGEHGIPVVVDSPGVGQNLQEHPGVLMSTHVGIDSLNVEVQSVARIVKHGLNFALFGRGPATACVASALAFIRTRDHLEWPNIQLSFSPIAYDFTPDGVHLYKRAAIGVAINICRPETRGQLLLRSTDPSERPIIQHELLGGDDEIKQLIEGCRIVRKIFRSKPFSEYDKGERLPGKQVETDADWIEYIRQSAFLMYHPTGTCAMGIGPTAVLDPELRVKGVTGLRVADASIMPTLVSANTNAPCIMIGERAADLIRRSH
